# BindCraft: one-shot design of functional protein binders

**DOI:** 10.1101/2024.09.30.615802

**Authors:** Martin Pacesa, Lennart Nickel, Christian Schellhaas, Joseph Schmidt, Ekaterina Pyatova, Lucas Kissling, Patrick Barendse, Jagrity Choudhury, Srajan Kapoor, Ana Alcaraz-Serna, Yehlin Cho, Kourosh H. Ghamary, Laura Vinué, Brahm J. Yachnin, Andrew M. Wollacott, Stephen Buckley, Adrie H. Westphal, Simon Lindhoud, Sandrine Georgeon, Casper A. Goverde, Georgios N. Hatzopoulos, Pierre Gönczy, Yannick D. Muller, Gerald Schwank, Daan C. Swarts, Alex J. Vecchio, Bernard L. Schneider, Sergey Ovchinnikov, Bruno E. Correia

**Author notes:** These authors contributed equally (Martin Pacesa, Lennart Nickel, Christian Schellhaas).

## Abstract

Protein–protein interactions (PPIs) are at the core of all key biological processes. However, the complexity of the structural features that determine PPIs makes their design challenging. We present BindCraft, an open-source and automated pipeline for *de novo* protein binder design with experimental success rates of 10-100%. BindCraft leverages the weights of AlphaFold2^1^ to generate binders with nanomolar affinity without the need for high-throughput screening or experimental optimization, even in the absence of known binding sites. We successfully designed binders against a diverse set of challenging targets, including cell-surface receptors, common allergens, *de novo* designed proteins, and multi-domain nucleases, such as CRISPR-Cas9. We showcase the functional and therapeutic potential of designed binders by reducing IgE binding to birch allergen in patient-derived samples, modulating Cas9 gene editing activity, and reducing the cytotoxicity of a foodborne bacterial enterotoxin. Lastly, we utilize cell surface receptor-specific binders to redirect AAV capsids for targeted gene delivery. This work represents a significant advancement towards a “one design-one binder” approach in computational design, with immense potential in therapeutics, diagnostics, and biotechnology.

## Introduction

Proteins are versatile biomolecules capable of mediating a diverse range of biological functions, including catalysis, molecular recognition, structural support and others. However, proteins rarely perform their biological functions in isolation but rather rely on protein–protein interactions (PPIs) to execute complex biological processes, such as signal transduction, antibody-mediated immunity, cellular communication, etc.

Designing protein binders that can specifically target and regulate PPIs holds immense therapeutic and biotechnological potential. Such binders can be utilized to modulate protein interaction networks and signaling pathways, design therapeutic antibodies, inhibit pathogenic agents, or create biotechnological tools for research and industry. However, traditional methods for generating protein binders, such as immunization, antibody library screening, or directed evolution of binding scaffolds, are often laborious, time-consuming, and provide limited control over the target site.

Computational protein design provides a powerful alternative, where binders can be designed and tailored to a specific protein target and binding site, enabling the exploration of a much broader sequence and structure space. For example, *de novo* designed protein binders have been previously used to block viral entry^2^, modulate immune and inflammatory response^3,4^, prevent amyloid assembly^5^, or control cell differentiation pathways^6^.

Physics-based protein design methods like Rosetta have been instrumental in early binder designs through scaffolding and sidechain optimization^7–9^. However, such methods suffer from very low experimental success rates (typically less than 0.1%) and require the generation and sampling of a vast number of designs, ranging from hundreds of thousands to millions^7,9–11^. Moreover, because such methods typically require the docking of predefined scaffolds onto a fixed target structure, incompatibilities between the target and binder surfaces can result in suboptimal binding interactions or even preclude the targeting of certain target sites.

Recent breakthroughs in deep learning have revolutionized the field of biomolecular modelling, particularly the prediction of protein structure. Models like AlphaFold2 (AF2)^1^ and RoseTTAFold2 (RF2)^12^, trained on large protein structure and sequence datasets, have demonstrated remarkable capabilities in accurately predicting protein structures and modeling complex PPIs. Indeed, AF2 filtering has been shown to significantly increase the success rates of binder design by evaluating the plausibility of predicted complexes^10,11^. Deep learning has also been successfully applied for *de novo* design of proteins and binders. The current state-of-the-art methods involve the use of RFdiffusion^10^ for backbone generation coupled with ProteinMPNN sequence generation^13,14^. When applied to binder design, this approach has successfully designed binders against a variety of protein targets, with improved success rates compared to previous methods^10^. However, it requires the generation of thousands to tens of thousands designs *in silico* and extensive experimental *in vitro* screening to identify suitable binders, which remains a significant limitation for most research groups.

Given the utility of AF2 in improving binder filtering success, we hypothesized that we could harness its weights and learned patterns of protein structures directly for the design of protein binders. We present BindCraft, a user-friendly pipeline for d*e novo* design of protein binders that requires minimal user intervention and computational expertise. BindCraft leverages backpropagation through the AF2 network to efficiently hallucinate novel binders and interfaces without the need for extensive sampling (**Fig. 1a**). We demonstrate the efficiency of our pipeline on twelve diverse, challenging, and therapeutically relevant protein targets (**Fig. 1b**) and identify several high affinity binders for each target without the need for high throughput screening of hundreds to thousands of designs experimentally. This marks an important advancement in the design of protein binders on demand, a long standing problem in protein design. Furthermore, BindCraft makes binder design accessible to research groups without means for high throughput screening facilities or expertise in computational design methods.

**Figure 1.**
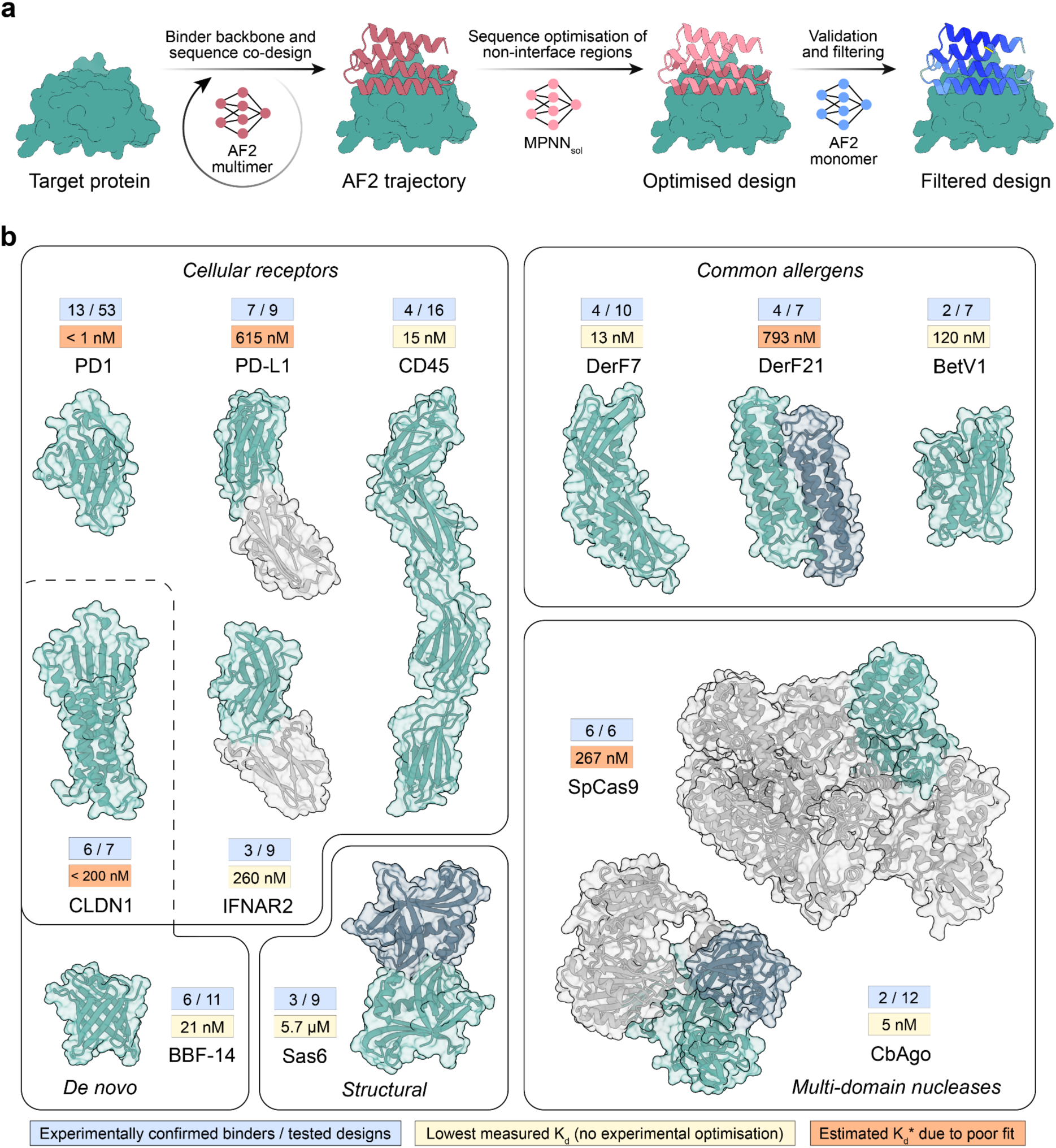
*De novo* binder design using BindCraft. **a,** Schematic representation of the BindCraft binder design pipeline. Given a target protein structure, a binder backbone and sequence is generated using AF2 multimer, then the surface and core of the binder are optimized using MPNN_sol_ while keeping the interface intact, and finally designs are filtered based on AF2 monomer model prediction. **b,** Overview of protein targets for binder design. Parts of the model colored in green were utilized during design, gray areas were excluded. Values in the blue box indicate the number of successful designs, where binding was observed on SPR measurement versus the total number of designs tested. Values in the yellow box indicate the measured K_d_ of the highest affinity binder without experimental sequence optimization, while values in orange boxes indicate estimated K_d_* values due to poor fit. PD1 binders were tested as a bivalent Fc-fusion.

## Results

### Deep learning-based design of *de novo* binders

Our goal was to create an accessible, efficient, and automated pipeline that leverages the capabilities of AF2 models for accurate binder design, reducing the need for large-scale sampling. To this end, the user inputs a structure of the target protein, the size range of binders, and the final number of desired designs. Target hotspots can be specified, or the AF2 network can automatically select an optimal binding site by identifying the one that satisfies the design criteria most effectively (see Methods). We utilize the ColabDesign implementation of AF2 to backpropagate hallucinated binder sequences through the trained AF2 weights and calculate an error gradient. This error gradient is used to update and optimize the binder sequence to fit specified loss functions and design criteria, similarly to previous protein hallucination approaches^15–18^. These include the AF2 confidence scores, intra- and intermolecular contacts, helical content, and radius of gyration. By iterating over the network, we can generate high quality designs, essentially enabling the generation of binder structure, sequence, and interface concurrently (**Fig. 1a**). Unlike other design methods, such as RFdiffusion^10^ or RIFdock^7,11^, which keep the target backbone fixed during design, BindCraft repredicts the structure of the binder-target complex at each design iteration. This allows for defined levels of flexibility on the side-chain and backbone level for both binder and target, resulting in backbones and interfaces that are molded to the target binding site. The root mean square deviation (RMSD_Cα_) of the target structure after binder design ranges from 0.5 - 5.5 Å, depending on the design target (**Extended Data Fig. 1a**). The target flexibility can further be increased by masking the sequence of the input template and providing only Cα coordinates, without affecting overall confidence of the complex prediction (**Extended Data Fig. 1b**). Additionally, to mitigate the generation of purely alpha-helical binders, we apply a “negative helicity loss”. This allows to increase the beta sheet content of our designs and even allow the generation of purely beta sheeted proteins (**Extended Data Fig. 1c**), although with reduced *in silico* design success rates (**Extended Data Fig. 1d-e**).

We utilize AF2 multimer^19^ for designing initial binders, as this version of AF2 was trained on protein complexes and would likely be able to more accurately model PPIs compared to AF2 monomer^1^. We note that AF2 multimer hallucinates on average 20% larger interfaces, with a larger proportion of loops, and higher confidence than the AF2 monomer model (**Extended Data Fig. 1f**). We utilize all 5 trained model weights of AF2 multimer to avoid overfitting of sequences to a single model. However, we and others^14,20^ have previously demonstrated that AF2-hallucinated proteins can exhibit low levels of expression when tested experimentally. We therefore subsequently optimize the sequence of the binder core and surface using MPNN_sol_^13,20^ while keeping the interface intact (**Fig. 1a**). The optimized binder sequences are repredicted using the AF2 monomer model^1^. This model was exclusively trained on monomeric proteins, which minimizes prediction bias of PPIs and enables robust filtering for high quality interfaces. Lastly, as deep learning models have been shown to sporadically produce physically improbable results^1,19^, we filter the predicted designs based on AF2 confidence metrics, as well as Rosetta physics-based scoring metrics (see Methods).

Each target exhibits varying levels of *in silico* design success, with 16.8 - 62.7% of initial AF2 trajectories displaying satisfactory confidence metrics, and 0.6 - 65.9% of MPNN_sol_-optimized designs passing the final computational filters after AF2 monomer complex reprediction (**Extended Data Fig. 1g**). Notably, design success rates are influenced by the size of the designed binder (**Extended Data Fig. 1h**). We demonstrate that BindCraft generates diverse backbone topologies optimized to geometrically complement the target site (**Extended Data Fig. 2a**) and enable the design of novel, high-affinity interfaces (**Extended Data Fig. 2b**).

All of the described steps are automated into a single workflow, where designs are filtered automatically, statistics are stored in a user-friendly format, and settings have been optimized to ensure the design procedure is generalizable across different targets. This allows research groups without protein design expertise to generate binders on demand for any application. By minimizing human intervention needed to generate and sort high quality binder designs, BindCraft democratizes protein binder design and makes it accessible to a broader scientific community.

### Targeting cell surface receptors

To test the performance of our pipeline, we designed binders against therapeutically-relevant cell surface receptors and tested them for binding activity *in vitro.* Receptors are ideal targets for binder design due to the presence of well-characterized binding sites exposed on the extracellular domain, either through interactions with endogenous binding partners or therapeutic antibodies. We first generated designs that could bind the human PD-1 protein, a key immune checkpoint receptor expressed primarily on the surface of T cells^21^. We initially purified and screened 53 designs for binding using bio-layer interferometry (BLI) in a bivalent Fc-fusion format. We observed a binding signal for 13 binders, with the best binder displaying an apparent dissociation constant (K_d_*) lower than 1 nM (**Fig. 2a-b**), although the exact K_d_ could not be determined due to the extremely slow dissociation rate and avidity effect from the Fc-fusion construct. To confirm the binding site, we performed a competition assay with the well characterized anti-PD-1 monoclonal antibody, pembrolizumab, which should engage the same binding site. Indeed, our binder could not outcompete the antibody binding (K_d_ = 27 pM), indicating it is targeting overlapping binding sites (**Fig. 2c**).

**Figure 2.**
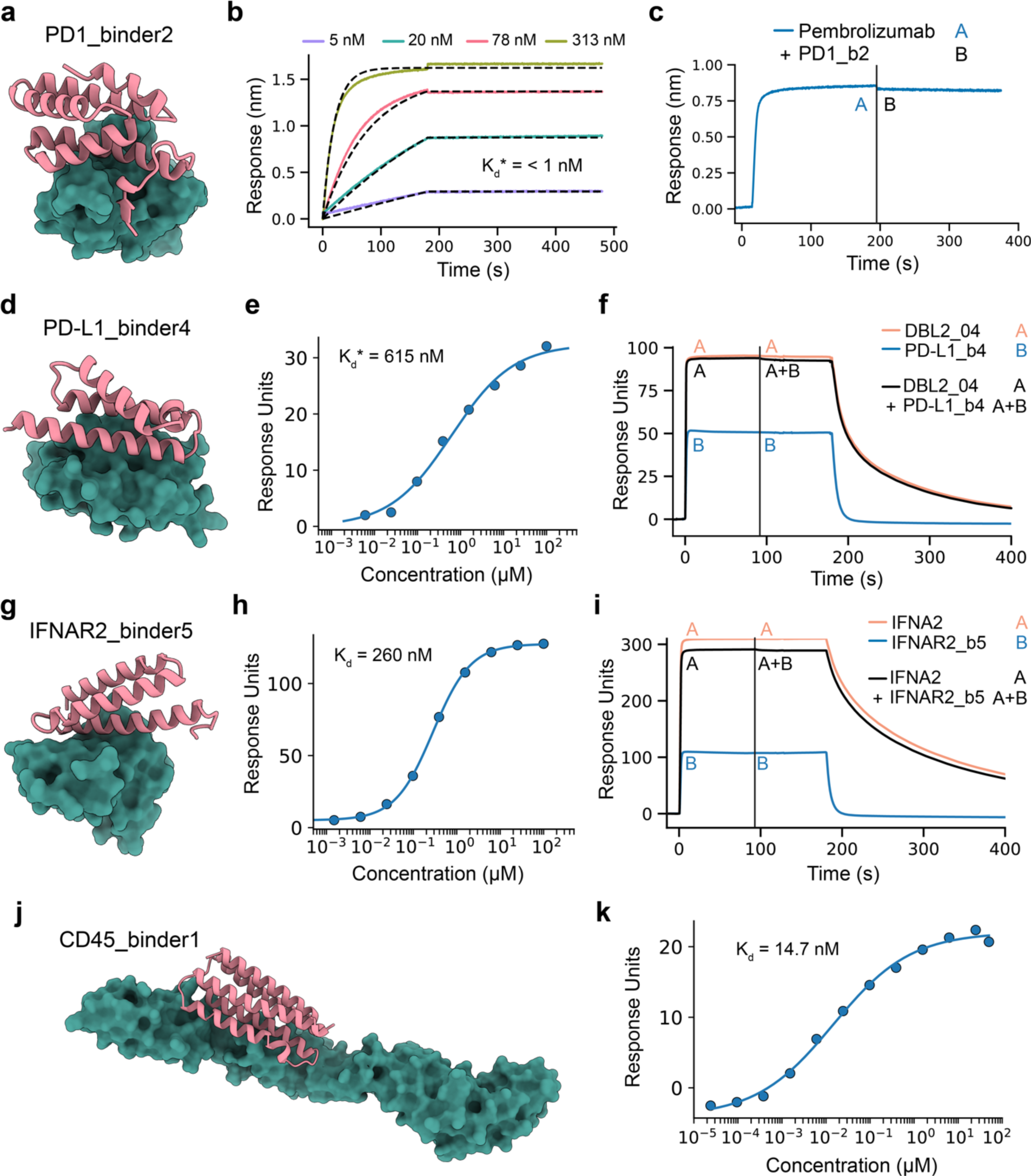
Binder design targeting cell surface receptors. **a,** Design model of binder2 in complex with PD-1. **b,** Representative BLI sensorgram showing binding kinetics of binder2 (bivalent Fc-fusion) to PD-1. **c,** Competition assay showing that the anti-PD-1 antibody, pembrolizumab, occupies the same binding site on PD1 as binder2. **d,** Design model of binder4 in complex with PD-L1. **e,** Binding affinity determination by SPR for the PD-L1-binder4 interaction. **f,** Competition assay showing previously published *de novo* binder DBL2_04 occupying the same binding site as PD-L1 binder4. **g,** Design model of binder5 in complex with IFNAR2. **h,** Binding affinity determination by SPR for the IFNAR2-binder5 interaction. **i,** Competition assay displaying the natural IFNA2 binding partner occupies an overlapping binding site with IFNAR2 binder5. **j,** Design model of binder1 in complex with CD45. **k,** SPR binding affinity fit for binder1.

Encouraged by these results we reduced the number of designs tested experimentally for all subsequent targets to test whether we can minimize the need for experimental screening. We next designed binders against PD-L1^21^ and the interferon 2 receptor (IFNAR2)^22^, both important modulators of immune signaling, where specific binders could enable the design of novel tumor or antiviral therapies. We tested 9 designs against PD-L1 out of which 7 showed a binding signal (**Extended Data Fig. 3a**), while for IFNAR2 we could detect binding for 3 out of 9 designs (**Fig. 1b**). The top performing binder4 against PD-L1 displayed an apparent K_d_* of 615 nM (**Fig. 2d-e**) and an expected alpha-helical signature as measured by circular dichroism (CD) (**Extended Data Fig. 3b**). We probed the binding of our PD-L1 binder4 using a previously characterized *de novo* designed binder^8^ and could confirm they compete for the intended target binding site (**Fig. 2f**). Interestingly, the designed binder engages in a distinct mode of binding compared to PD1, while partially occupying the same binding site (**Extended Data Fig. 3c**).

The top performing binder5 against IFNAR2 displayed an affinity of 260 nM determined by surface plasmon resonance (SPR) (**Fig. 2g-h**) and a typical alpha-helical signature by CD, validating the fold integrity and the thermal stability of our designs (**Extended Data Fig. 3d**). Similarly to PD-L1, we probed the binding of IFNAR2 binder5 against its native binding partner, the cytokine interferon alpha 2 (IFNA2)^22^. We observe competition for the native IFNA2 binding site, validating our designed binding mode (**Fig. 2i**). Notably, binder5 and the native IFNA2 binding site exhibit only minimal overlap, primarily occupying distinct binding sites (**Extended Data Fig. 3e**). These results demonstrate we are able to efficiently design binders, straight from the computational design pipeline, against known binding sites, without the need for extensive screening to identify hits with nanomolar affinity.

Next, we sought to determine whether our pipeline could design binders against extracellular receptors lacking well-characterized binding sites. We selected CD45 as a target, due to the structural complexity of its extracellular domain (ECD), comprising four immunoglobulin (IG)-like domains d1-d4 (**Fig. 2j**) with heavy N-glycosylation in the smallest isoform^23^. CD45 is a transmembrane tyrosine phosphatase involved in critical pathways of T cell function. Designing binders against the ECD of CD45 could allow us to modulate its signaling activity and fine tune T-cell activation thresholds for anti-tumor therapies. As the ECD is large, we designed binders targeting individual domains and pairs of adjacent domains, while manually removing any binders that overlapped with known glycosylation sites. We tested 16 binders experimentally out of which 4 showed binding on SPR (**Fig. 1b**). The best performing binder1 showed a Kd of 14.7 nM and targeted the junction region of domains d3 and d4 (**Fig. 2j-k**). We also observed the expected alpha-helical signal in CD, validating the correct folding of our design (**Extended Data Fig. 3f**). These results indicate that BindCraft can effectively design binders against novel or previously uncharacterized binding sites.

### Binder design beyond known and natural binding sites

Membrane proteins lacking distinct extracellular domains are of critical biological and therapeutic importance; however, they pose a significant challenge for binder design due to difficulties in experimental validation and screening. The use of computationally designed soluble analogues that retain natural epitopes^20^ offers a promising solution by enabling rapid prescreening of potential binders. We aimed to validate this approach by targeting claudins, membrane proteins critical for maintaining epithelial and endothelial tight junction barrier integrity^24^. Dysregulation of claudins contributes to diseases such as inflammatory disorders, cancer, and microbial infections^24^. Claudins are naturally targeted by *Clostridium perfringens* enterotoxin (CpE), which binds their extracellular domain, to form a membrane-penetrating pore that leads to cell death^25^. We hypothesized that binders could be designed to bind claudins and compete with CpE for its binding site to mitigate CpEs cytotoxicity (**Fig. 3a**).

**Figure 3.**
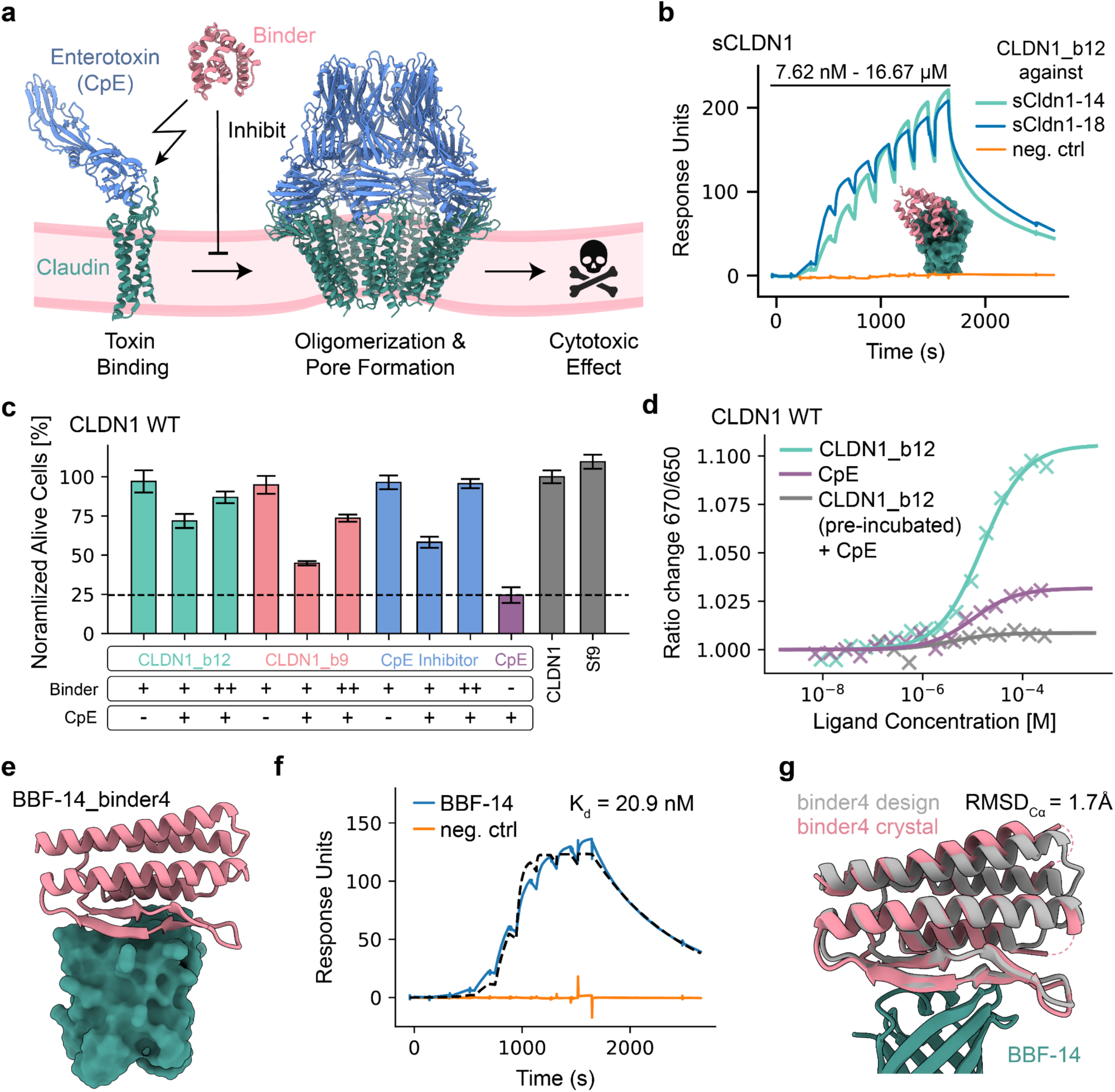
Targeting natural and *de novo* binding epitopes. **a,** Schematic of CpE-based cytotoxicity and CLDN1 binder inhibition. **b,** Single cycle kinetic analysis with SPR of CLDN1 binder12 binding to soluble analogues of CLDN1. **c,** Cell-based assay showing concentration dependent inhibition of CpE cytotoxicity by binder9, binder12 and CpE Inhibitor. **d,** MST measurements showing blocking of CpE binding to CLDN1 wild type when pre-incubated with binder12. **e,** Design model of binder4 in complex with *de novo* designed beta barrel BBF-14. **f,** Single cycle kinetic analysis with SPR of binder4 binding to BBF-14. g, Comparison of crystal structure (colored) of the BBF-14_binder4 complex overlaid with the design model (gray).

Using a soluble analogue of claudin 1 (sCLDN1) as a target^20^, we designed binders against claudin’s extracellular domain and prescreened them for binding using two variants of the soluble analogue (**Extended Data Fig. 4a-b**). We tested seven binders and found all except binder17 bound to sCLDN1-14 and sCLDN1-18 (**Extended Data Fig. 4a**), which both harbor the native CLDN1 extracellular epitope (**Extended Data Fig. 4b**). We observed the strongest binding signal for binder12, which exhibited nanomolar affinity for the soluble analogues (**Fig. 3b**). To assess the binder’s utility, they were tested further against wild-type claudin 1 (CLDN1 WT) using a cell-based cytotoxicity assay. Here, binder9 and binder12 effectively inhibited CpE-based cytotoxicity, protecting CLDN1 WT-expressing cells from cell death in a concentration-dependent manner and on the order of a known CpE inhibitor (**Fig. 3c, Extended Data Fig. 4c**). Notably, both of these binders result from the same initial trajectory and carry the same interface residues. To validate that the inhibition of cytotoxicity by the designs was the result of direct interactions with CLDN1 WT, we used microscale thermophoresis (MST) and found that both CpE and binder12 interacted with CLDN1 WT, and that preincubation of binder12 with CLDN1 WT blocked CpE binding, indicating competition for the same extracellular binding site (**Fig. 3d**). Interestingly, the binders failed to protect claudin 4 (CLDN4)-expressing cells from CpE-induced toxicity (**Extended Data Fig. 4d-e**). This is most likely due to differences in binding mode (**Extended Data Fig. 4f**) and CpE’s ∼400-fold higher affinity for CLDN4^25^, which might outcompete the lower-affinity binders. Overall, these results suggest that designing binders against soluble analogues of membrane proteins holds promise in discovering binders that are useful in modulating the functions of natural membrane proteins, given the accuracy of the soluble form.

To further assess the generalizability of our pipeline for targeting proteins or surfaces lacking known binding sites, we designed binders against a protein with no known sequence homologs in the PDB. We chose the *de novo* designed beta-barrel fold 14 (BBF-14) as a target^20^, as beta barrels are not commonly regarded as PPI partners. We purified the 11 top-scoring designs from which 6 showed binding (**Fig. 1b**). The best binder, binder4 (**Fig. 3e**), is composed of a mixed alpha-beta topology, with the interface formed by both the split beta-sheets and a helix motif. Interestingly, the beta-sheet interface is not mediated by backbone hydrogen bonding, but rather by sidechain interactions. Binder4 exhibited a Kd of 20.9 nM for BBF-14, as determined by SPR (**Fig. 3f**). To assess the fidelity of our design procedure, we solved a structure of BBF-14 bound to binder4 (**Extended Data Fig. 5a**). When aligned on the BBF-14 target, binder4 exhibited a backbone RMSD_Cα_ of 1.7 Å, confirming both the accuracy of the fold and the designed binding mode (**Fig. 3g**). This result underscores our ability to generate binders purely based on structural information, without relying on existing binding sites or any influence from co-evolutionary data.

We additionally chose the highly conserved structural protein SAS-6 as a design target. SAS-6 assembles into higher-order oligomers and is essential for centriole biogenesis across the eukaryotic tree of life^26^. A major challenge in the study of the centriole’s architecture has been the lack of tools that allow precise modulation of the assembly process. We attempted binder design against *Chlamydomonas reinhardtii* SAS-6 previously using published computational methods^8^ but were unable to obtain satisfactory binders. Using BindCraft, we generated several designs passing computational filters, and experimentally tested 9 top scoring designs. We identified binder4 (**Extended Data Fig. 5b**) that binds with 5.7 μM affinity to the monomeric form of CrSAS-6 (**Extended Data Fig. 5c**) and 4.2 μM affinity to the dimeric form (**Extended Data Fig. 5d**), indicating compatibility with its oligomeric form. Binder4 targets an overlapping target site with the previously reported monobody MB_CRS6_-15 (**Extended Data Fig. 5e**), which binds to the N-terminal head domain of CrSAS-6 and causes a shift in its assembly mechanism, transforming the ring-like structure into a helical assembly^27^. We speculate that we can now design binders on-demand against challenging targets to probe their biological function, even in the context of higher order assemblies.

### Blocking immunogenic epitopes of common allergens

The prevalence of allergic rhinitis and seasonal allergies have been estimated to affect up to 50% of the population in some countries^28^. Current treatments primarily focus on reducing global inflammation with immunosuppressants and monoclonal antibodies. However, neutralizing allergic reactions could potentially offer a more effective strategy for managing allergies. Allergens comprise a diverse group of proteins with different folds, biological functions, and highly charged surfaces^29^. Generally, hydrophobic binding sites are considered more tractable for computational binder design^7^, making allergens more challenging targets.

To test the capabilities of BindCraft at targeting allergens, we designed binders against the dust mite allergens Der f7 and Der f21, and the major birch allergen Bet v1, which is responsible for up to 95% of birch-related allergies^30^. We examined 10 designs against Der f7 experimentally and identified 4 binders (**Fig. 1b**), with binder2 exhibiting the highest binding affinity with a K_d_ of 12.8 nM (**Fig. 4a**). To confirm the binding mode of binder2, we solved crystal structures in complex with Der f7 obtaining two crystal forms with resolutions of 2.2 Å and 3.0 Å (**Extended Data Fig. 6a-b, Extended Data Table 1**). When aligned on the allergen, the backbone RMSD_Cα_ of binder2 compared to the design model is 1.7 Å (**Fig. 4b**), validating the structural accuracy of the design method. Interestingly, binder2 exhibits a helical topology that wraps around connecting loops of the mixed beta sheet and helical tip of the protein (**Fig. 4b**). Mouse monoclonal antibodies raised against Der f7 have been shown to bind to the same epitope through mutational studies^31^, indicating that we are able to target known immunogenic epitopes of allergens.

**Figure 4.**
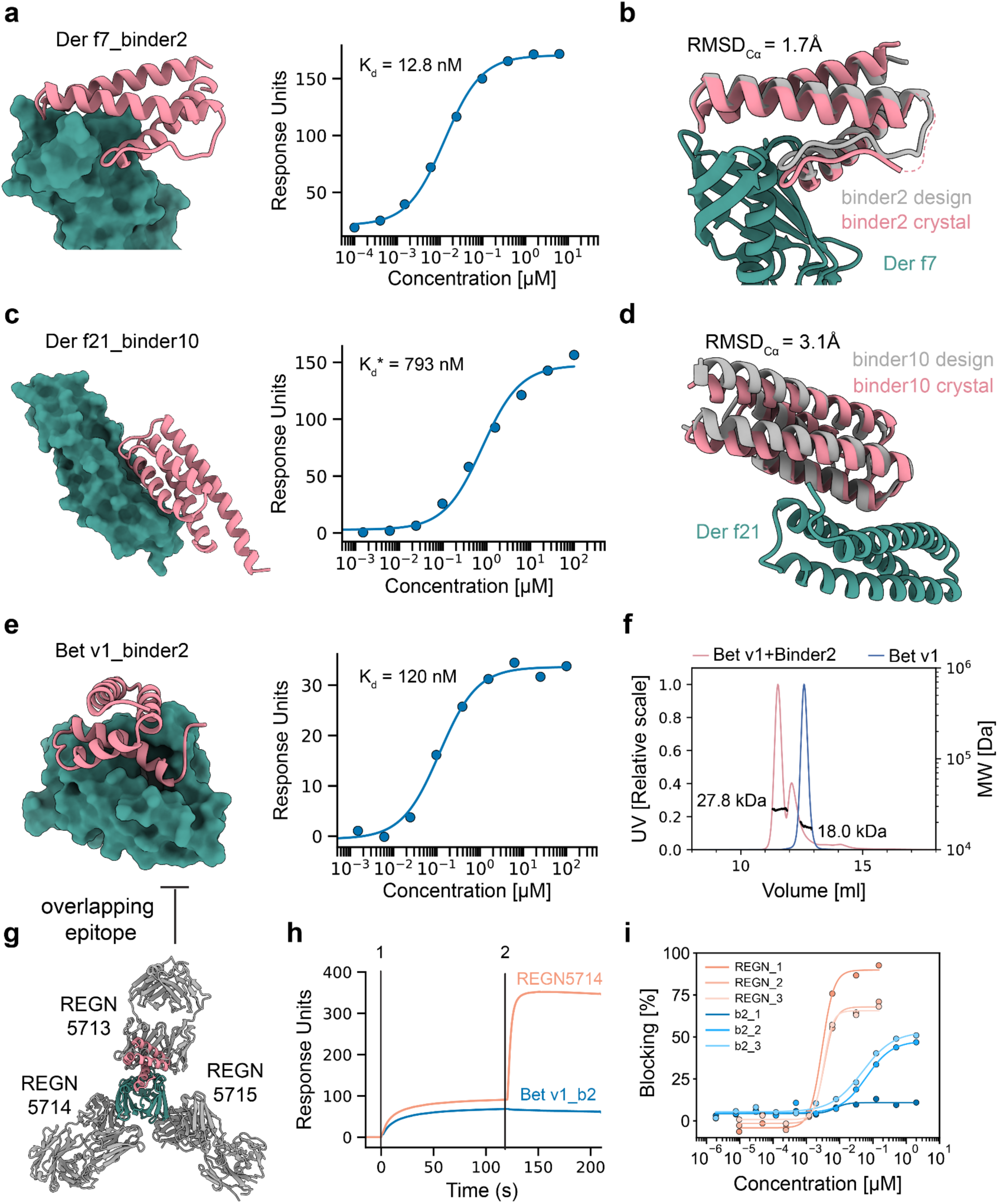
Designs occluding epitopes of common allergens. **a,** Left: Design model of binder2 against dust mite allergen Der f7. Right: SPR binding affinity fit for binder2. **b,** Crystal structure (colored) of the Der f7_binder2 complex overlaid with the design model (gray). **c,** Left: Design model of binder10 against dust mite allergen Der f21. Right: SPR binding affinity fit for binder10. **d,** Crystal structure (colored) of the DerF21_binder10 complex overlaid with the design model (gray). **e,** Left: Design model of binder2 against birch allergen Bet v1. Right: SPR binding affinity fit for binder2. **f,** SEC-MALS analysis of Bet v1 allergen (blue) and Bet v1 mixed with binder2 (orange). **g,** CryoEM structure (PDB: 7MXL) of Bet v1 bound to commercial anti-Bet v1 REGN antibody mix. **h,** Competition assay on immobilized REGN5713-Bet v1 complex binding of the REGN5714 antibody but not Bet v1 binder2, confirming binding at the designed site. **i,** Blocking ELISA showing the capacity of the REGN antibody mix (orange) or binder2 (blue) to prevent the binding of Bet v1 to IgE from three birch-allergic patient sera. Number suffix represents individual serum from a patient.

Similarly, we evaluated 7 binders against Der f21 and could detect binding for 4 designs by SPR (**Fig. 1b**). The best performing binder10 displayed an apparent affinity of 793 nM (**Fig. 4c**). We solved a 2.6 Å resolution crystal structure to validate the mode of binding of binder10 against a highly charged helical site of Der f21 (**Extended Data Fig. 6c**), the structure and the predicted model differed in terms of backbone RMSD_Cα_ by 3.1 Å, caused by an alternative rotamer conformation of an interface tyrosine (**Fig. 4d**). Similarly to Der f7, structures of Der f21 bound to induced antibodies are not available, however mutational analysis in patient sera indicated that our binders target epitopes distinct from the IgE sera of allergic individuals^32^.

Lastly, we tested 7 binders against the birch allergen Bet v1 and could identify 2 successful binders (**Fig. 1b**). Binder2 exhibited a 120 nM binding affinity by SPR (**Fig. 4e**) and we could further validate the binding interactions using size exclusion chromatography with multi-angle light scattering (SEC-MALS) where the complex shows the expected mass of 27.8 kDa (**Fig. 4f**). The binder2 exhibits a warped helical topology, where its C-terminal helix inserts itself deep into the ligand binding pocket of Bet v1^33^. Previously, an antibody cocktail mix of three antibodies that bind three different immunogenic epitopes of Bet v1 was developed to prevent allergic response^34^.

The published cryoEM structure indicates that our binder targets a known epitope recognized by the REGN5713 antibody, albeit with a different binding mode (**Fig. 4g**). To probe the binding mode, we immobilized REGN5713 on SPR and loaded the Bet v1 allergen on it. We observe a binding signal with REGN5714 as the analyte, but not with binder2, confirming that it targets an overlapping epitope with REGN5713 (**Fig. 4h**). We further hypothesized that our binders can compete with Bet v1 specific IgE present in serum samples from birch allergic patients, similarly to the REGN antibody mix^34^. To test the neutralization activity of our anti-Bet v1 binder2, we performed a blocking ELISA using the serum of three birch allergic patients with high titers of anti-Bet v1 IgE. In this assay, biotinylated Bet v1 was preincubated with either the REGN antibody cocktail or our designed binder2 (**Fig. 4i**). While the REGN three antibody mix was able to block up to 90% of the binding of Bet v1 to IgE at low concentrations, our single binder exhibited blocking rates of up to 50% in 2 out of 3 donors. This is on par with blocking rates of single antibodies^34^, indicating that there is therapeutic potential for *de novo* designed binders in neutralizing allergic responses targeting multiple epitopes.

### Modulating the function of large multi-domain nucleases

Nucleic acid interaction interfaces in proteins have long been considered to be undruggable^35^. This is due to their highly charged, convex and large interfaces, which are difficult to target with small molecules^35^. Protein binders offer a promising alternative to modulate protein-nucleic acid interactions in biotechnological and therapeutic applications. We decided to test the applicability of our pipeline to such interfaces on the large multi-domain CRISPR-Cas9 nuclease from *Streptococcus pyogenes* (SpCas9). SpCas9 has been adapted for gene editing applications due to its easy programmability and has since revolutionized synthetic biology and medicine^36^. However, CRISPR-Cas9 is originally a prokaryotic immune system that protects bacteria against invading genetic elements^37^. To counter this, phages have evolved small proteins, termed anti-CRISPRs (Acrs), that can block CRISPR-Cas nuclease activity by directly occluding nucleic acid binding sites^38^. We wondered whether we could design artificial Acrs that would emulate a similar function.

We designed binders against the bipartite REC1 domain of SpCas9, which contains a highly charged pocket for the binding of the guide RNA^39^ (**Fig. 5a**). We tested 6 binders experimentally and strikingly all 6 binders bound the full length apo SpCas9 enzyme (**Extended Data Fig. 7a**). The top performing binder3 and 10 exhibited apparent binding affinities in the range of 300 nM by SPR, although complete titration curves were experimentally challenging to obtain and therefore the real affinity might differ. To validate their binding mode, we attempted to solve cryoEM structures of binder3 and binder10 bound to the full length SpCas9 apo enzyme. Despite the high quality of the data in both cases, we were unable to obtain a satisfactory cryoEM density to build an atomic model (**Extended Data Fig. 7b-c**), presumably due to the dynamic nature of the apo form of Cas9^40^. Nevertheless, we observe clear density in the binding site of the REC1 domain and can confidently dock both binders, validating the designed binding mode (**Fig. 5b-c, Extended Data Fig. 7d-e**).

**Figure 5.**
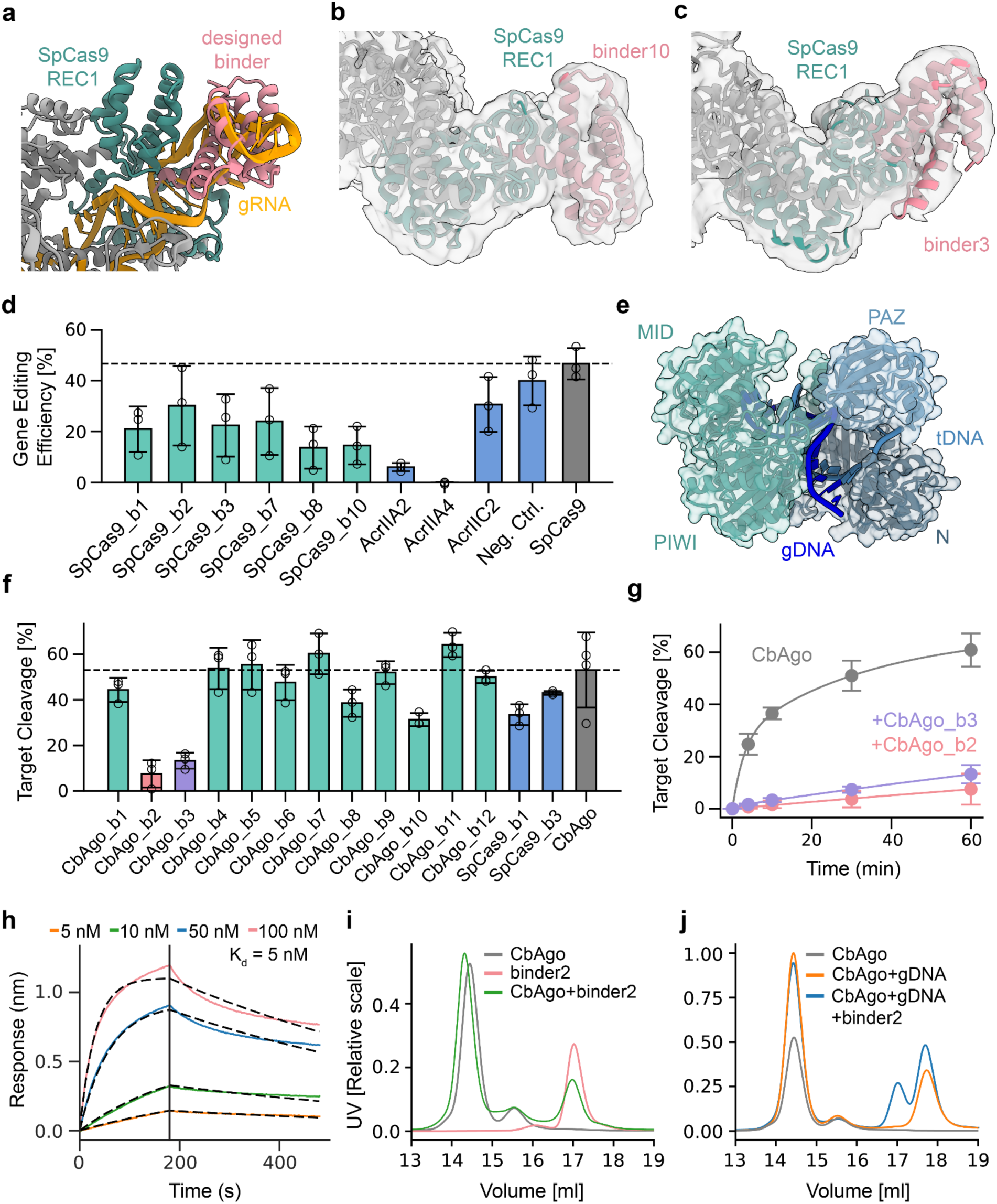
Targeting nucleic acid interactions with *de novo* binders against nucleic acid-guided multidomain nucleases. **a,** Zoom in on the SpCas9 REC1 domain with bound guide RNA (PDB: 4ZT0). A designed binder is overlaid in the binding pocket. **b,** CryoEM structure of binder3 bound to the apo form of SpCas9. REC1 domain is highlighted in green, rest of SpCas9 in gray. CryoEM density overlaid in gray. **c,** CryoEM structure of binder10 bound to the apo form of SpCas9. REC1 domain is highlighted in green, rest of SpCas9 colored in gray. CryoEM density overlaid in gray. **d,** SpCas9-based editing of HEK293T cells in absence (gray bar, dashed line) or in presence of designed binders (green bars) or natural Acrs (blue bars). **e**, Structural architecture of *Clostidium butyricum* Argonaute with bound gDNA and tDNA (PDB: 6QZK). PAZ domain and N+PIWI domains used as design targets are highlighted in light and dark blue. **f**, CbAgo-gDNA-mediated cleavage of target DNA in absence (gray bar, dashed line) or in presence of designed binders (green bars) or designed SpCas9 binders (blue bars). **g**, CbAgo-gDNA-mediated cleavage of target DNA in absence of binders (gray line) or in presence of designed binder2 (pink line) or binder3 (purple line). **h,** Representative BLI sensorgram displaying binding kinetics of CbAgo and binder2. **i**, Size exclusion chromatography (SEC) analysis of CbAgo only (gray line) or binder2 only (orange line) or combined (green line). **j**, SEC analysis of CbAgo only (gray line) or in presence of gDNA (orange line) or in presence of both gDNA and binder2 (blue line).

To evaluate the functional consequence of this binding, we co-transfected HEK293T cells with CRISPR-SpCas9 and either our designed binders or natural anti-CRISPR proteins^41–43^. We observe a significant reduction of SpCas9 gene editing activity in the presence of our designed binders (**Fig. 5d**). They outperform the natural AcrIIC2, which has also been shown to be an inhibitor of guide RNA loading, albeit using a different targeting mechanism^42^. AcrIIA2 and AcrIIA4, which inhibit target DNA binding (**Extended Data Fig. 7f**), nearly eliminate gene editing activity, underscoring the differences in inhibitory strategies. These results demonstrate that BindCraft-generated binders induced novel modes of nuclease inhibition that are distinct from known natural mechanisms.

To expand our binder design to other large nucleases, we designed binders against the multi-domain Argonaute (Ago) nuclease from *Clostridium butyricum* (CbAgo). The programmable nature of CbAgo and related prokaryotic Agos has been exploited for the development of numerous diagnostics and *in vivo* biotechnological applications^44^. Akin to Cas9, CbAgo acts as an immune system that utilizes a small oligonucleotide guide to target and cleave invading DNA^45,46^. However, in contrast to Cas9, CbAgo utilizes a small guide DNA (gDNA) to target single stranded target DNA (tDNA), and to date no natural inhibitors of Argonaute nucleases have been described. We designed binders targeting either the N-PIWI channel or the PAZ domain of CbAgo (**Fig. 5e**). We tested the effect of 12 binders on CbAgo-gDNA-mediated tDNA cleavage and two binders strongly inhibit CbAgo activity (**Fig. 5f**). Whereas CbAgo alone has a k_cat_ of 0.004 s^-1^, in presence of binder2 and binder3 the k_cat_ is reduced 80-fold to 5*10^-5^ s^-1^ and 40-fold to 9.8*10^-5^ s^-1^, respectively (**Fig. 5g**). We found that binder2 binds to CbAgo with a K_d_ of 5 nM, as determined by BLI (**Fig. 5h**). Size exclusion chromatography analysis of binder2 with CbAgo validates that it forms a stable complex with CbAgo (**Fig. 5i**). Addition of the gDNA destabilizes the CbAgo-binder2 complex, which confirms that binder2 occupies the gDNA binding channel (**Fig. 5j, Extended Data Fig. 7g**).

Combined, the experimental validation of our designed SpCas9 and CbAgo binders demonstrates that we can design protein binders even against challenging nucleic acid binding sites and grooves, potentially opening paths towards novel types of protein-based therapeutics and gene editing modulators. In addition, computational protein design now enables the design of inhibitors and modulators of molecular systems for which natural analogues have yet to be discovered, opening many possibilities for the generation of novel molecular biology tools for basic research.

### AAV re-targeting for cell-type specific gene delivery

Viral vectors, such as those derived from adeno-associated viruses (AAVs), have expanded gene therapy possibilities by leveraging the natural ability of viruses to introduce genetic material into cells and tissues. However, AAVs have poor specificity to cell types, tissues and organs. Targeting the cells of interest therefore often necessitates high vector doses, which increases the risk of off-target effects. Several efforts have been made to modify the tropism of AAV capsids towards more specific transduction profiles, by insertion of peptide segments ^47,48^ or receptor-binding moieties, such as nanobodies or DARPins^49,50^. However, such approaches involve library screening or immunization campaigns, usually with limited control over the target site. We hypothesized that BindCraft could efficiently design miniprotein binders capable of re-targeting AAVs to cell-type specific receptors (**Fig. 6a**). In addition, the high experimental success rates could allow the direct testing of AAV transduction *in cellulo*, without biochemical pre-screening of the designed binders. This would provide a platform for the rapid development of re-targeted AAV vectors to cells and tissues of interest.

**Figure 6.**
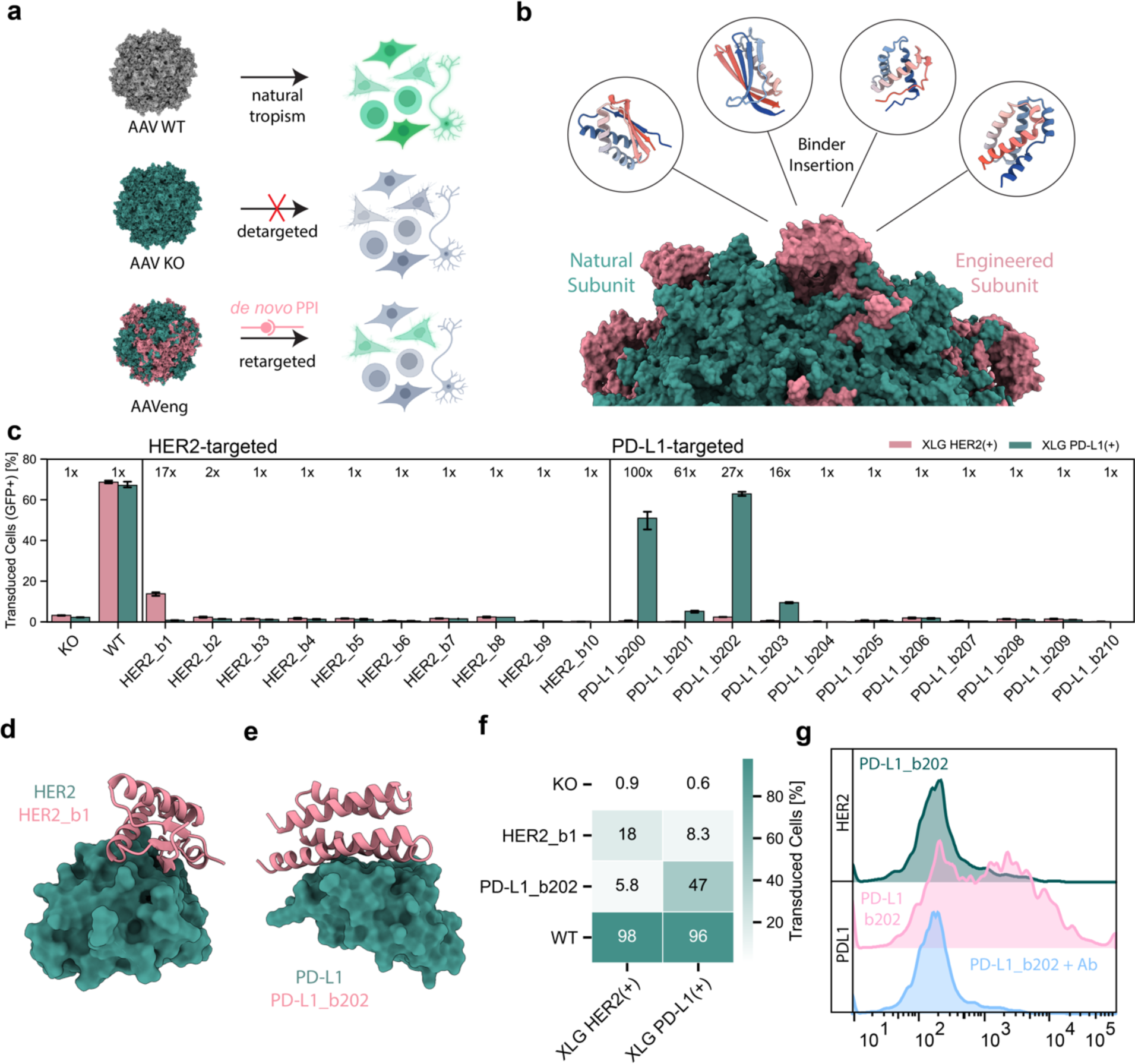
Engineering targeted gene delivery via AAV. **a,** Schematic representation illustrating AAV-cmv-GFP re-targeting upon genetic insertion of a cell-type receptor specific miniprotein binder, replacing the natural primary attachment to cell-surface glycans. **b,** Chimeric assembly of a re-targeted AAV particle, composed of the capsid proteins with (pink) and without (green) inserted binder in a defined stoichiometric ratio. **c,** Transduction efficiency measured by flow cytometry of different AAV variants targeting HER2 or PD-L1, determined after transfer of packaging cell supernatant onto HEK293 cells stably overexpressing the respective target receptors. The signal-to-noise ratio, defined as target/non-target ratio between the transduction rates measured on each cell line, is indicated as ‘x’ fold change. For comparison, each of the two cell lines is similarly transduced with the wild-type AAV6-cmv-GFP (WT) and the AAV capsid variant carrying knock-out mutations (KO). **d,** Design model of binder1 against HER2. **e,** Design model of binder202 against PD-L1. **f,** Heatmap of the transduction rates at a normalized multiplicity of infection (MOI) of 1*10^5^ vg/cell of the AAV variants carrying the binder1 against HER2 and binder202 against PD-L1, as well as the KO and WT controls, on HEK293 cells stably overexpressing the respective target receptors. **g,** Transduction with the PD-L1-targeting AAV carrying the binder202. The lower histogram shows that an anti PD-L1 antibody, that targets the binding site of AAV-binder202, blocks the transduction of HEK293 cells stably overexpressing PD-L1.

Traditionally, re-targeting molecules are either inserted into the variable regions VR-IV or VR-VIII, protruding near the three-fold symmetry axis of the AAV capsid, or fused to the N-terminus of the viral capsid protein 2 (VP2). Based on a large mutational study on AAV capsid fitness^51^, we explored an alternative insertion site, located between residues 497 and 498 of the VR-V near the three-fold symmetry axis of the AAV capsid (**Fig. 6b**). We chose AAV6-cmv-GFP as a starting vector and introduced point mutations to deplete its natural primary interactions with heparin and sialic acid (KO, **Fig. 6a**). We then designed binders against HER2 and PDL1 with an additional NC termini distance loss to facilitate a direct capsid integration, using a short -(GSG)_1_- extension on each terminus (**Fig. 6b**). To simultaneously screen the designed AAVs for production and transduction efficiency, a small-scale assay was designed that relies on directly transferring the supernatant of AAV-packaging cells onto the targeted cells (**Extended Data Fig. 8a-b**). This assay led us to identify one reprogrammed AAV to target HER2 and four targeting PD-L1 that showed enhanced specificity for cells (HEK293) stably overexpressing their respective target receptor (**Fig. 6c, Extended Data Fig. 8c**). We selected two variants, HER2_b1 and PD-L1_b202 (**Fig. 6d-e**), based on their transduction efficiency, for further characterization and showed that both AAVs had enhanced specificity towards cells expressing their target receptor (**Fig. 6f**). When the interaction was challenged with an antibody targeting the same receptor-binding site, the transduction of PD-L1-expressing cells by the PD-L1-targeting AAV was blocked, suggesting that the designed binder mediates the transduction through the engagement with the target receptor (**Fig. 6g**).

## Discussion

The design of *de novo* PPIs by computational means has been a cornerstone problem in protein design. This is primarily due to our lack of detailed understanding of the determinants of molecular recognition that drive PPIs and protein-ligand interactions. Recent advances in deep learning, particularly the development of accurate structure prediction networks such as AF2^1,19^, have revolutionized the field and enabled more accurate filtering of *de novo* designs with favorable biochemical profiles. Here we introduce a robust pipeline based on backpropagation through the AF2 network, an approach that has been explored in previous studies^15–18,52^, and extending its capabilities to the hallucination of protein binders. Unlike the majority of previously described approaches, BindCraft allows for flexibility on the target protein, which given the intrinsic flexibility of protein structures could be critical for capturing binding-induced changes essential for effective molecular recognition.

Our results demonstrate the performance of BindCraft in designing binders against a diverse set of 12 challenging targets. The binder affinities lie predominantly in the nanomolar range, with one in the micromolar range. The success rates range from 10 to 100%, with an average success rate of 46.3%, which is remarkable for designs resulting from a purely computational approach. These rates allow for the screening of far fewer designs experimentally to identify functioning binders, when compared to the current state of the art RFdiffusion^10^ and the recently described closed-source AlphaProteo binder design pipeline^53^. Notably, a binder designed with our pipeline recently ranked first in a community-wide binder design competition, showing nanomolar affinity against the challenging EGFR target^54^. However, we do expect success rates to vary based on the target protein and desired binding site.

One of the main challenges of PPI design is the choice of a favorable target site^7,11,55^. Prototypical binding sites are often composed of hydrophobic patches with mostly flat surfaces. Here we targeted a wide range of sites with diverse features, some with previously described binding interfaces, such as in the case of cell surface receptors, as well as unexplored surface regions in *de novo* proteins, allergens, and CRISPR-Cas nucleases. In the absence of a defined target site, BindCraft is able to sample optimal binding sites by making use of the trained AF2 multimer model^19^, suggesting that the network has likely learned which sites have a high propensity for forming PPIs. This is the case even for challenging binding sites, such as protein-nucleic acid interfaces, which will potentially unlock new avenues for the design of transcription factor modulators, whose aberrant activity is the underlying cause of many oncogenic diseases^35^.

The structural accuracy of our method, validated through both crystallography and cryoEM, not only allows us to create proteins that bind to defined surfaces but also enables their application for biotechnological and therapeutic advancements. We demonstrate this by utilizing our designed binders to reduce the binding of birch allergen Bet v1 to specific IgE from patient-derived serum samples. While a single binder displayed limited blocking activity compared to an antibody cocktail, we anticipate that covering a larger part of the antigen surface could produce comparable results. *De novo* binders would therefore offer a promising alternative to antibodies for such treatments, due to their high stability. However, due to the synthetic nature of our binders and their relatively large size (60–240 amino acids), concerns about immunogenicity and effective delivery persist, though these issues are gradually being addressed in preclinical models^3^.

BindCrafts’s high experimental success rates allow direct screening of intended biological function, without biochemical pre-screening, as exemplified by the re-targeting of AAV towards specific cell-surface receptors. The approach demonstrates the potential of computationally designed binders to efficiently re-target AAVs, enabling precise and customizable transduction profiles. Overall, the reported strategy promises to simplify the development of targeted viral vectors, offering a versatile platform for gene therapy applications, including therapeutic delivery to disease-relevant cells and tissues while minimizing the risk of potential off-target effects.

Despite the design successes outlined here, there are limitations to the BindCraft design approach. Backpropagation through the AF2 network requires the use of a GPU with large amounts of memory. For instance, a target-binder complex 500 amino acids in size allocates about 30 Gb of GPU memory. This sometimes requires the trimming or splitting of large proteins and complexes during binder design. Additionally, since we utilize AF2 monomer in single sequence mode for filtering, it is possible that we exclude prospective high affinity binders at the cost of a robust binding predictor. This is exemplified by AF2 monomer not being capable of predicting several experimentally validated high affinity complexes and can still predict false positives^7,8,10,11,56^ (**Extended Data Fig. 9a-b**).

AF2 has also been shown to be insensitive to the predicted effects of point mutations^57^, which could be detrimental at PPI interfaces, where a single mutation can abrogate or significantly enhance binding. The addition of an orthogonal physics-based scoring method, such as Rosetta, has been shown to add more discriminatory power to binder identification^58^. Lastly, a potential limitation is the use of the AF2 i_pTM metric for the ranking of designs, which has emerged as a powerful binary predictor of binding activity (**Extended Data Fig. 9a-b**), but does not correlate with the interaction affinity^59^. However, accurate *in silico* prediction of affinity remains highly challenging and alternative ranking metrics may similarly struggle to address this complexity.

Looking forward, we aim to target even more challenging and diverse targets and diversify the structure of our designs towards more natural and complex folds, such as using conditioning towards specific therapeutically applicable protein families. Lastly, as none of the current *in silico* metrics have been shown to be predictive of affinity, there is the need to develop novel scoring approaches for ranking the quality of binders. Nevertheless, BindCraft represents a significant leap in the accurate design of binders for direct functional applications. We envision that through iterative refinement of our pipeline, we will eventually reach a ‘one design, one binder’ stage, omitting the need for screening. This will enable rapid generation of binders for applications in research, biotechnology, and therapeutics for a wide range of research groups without protein design expertise.

## Code availability

The full BindCraft code along with installation instructions and binder design protocols are available on GitHub under MIT license (https://github.com/martinpacesa/BindCraft). A Google Colab notebook for running BindCraft is available at https://github.com/martinpacesa/BindCraft/blob/main/notebooks/BindCraft.ipynb.

## Data availability

Atomic coordinates and structure factors of the reported X-ray structures have been deposited in the PDB under accession numbers 9HAC (BBF-14_b4), 9HAD (DerF21_b10), 9HAE (DerF7_b2, crystal form P21), and 9HAF (DerF7_b2, crystal form C121). CryoEM maps will be deposited in the Electron Microscopy Data Bank following further refinement. Structural models of binders are available at https://doi.org/10.5281/zenodo.14249738.

## Acknowledgments

We thank SCITAS at EPFL for support in running design trajectories. We thank Anthony Marchand, Petra E.M. Balbi, Ahmed Sadek, and Simon Mauro for support with protein purification. We thank Florence Pojer, Kelvin Lau, and Amedé Larabi (Protein Production and Structure Characterization Core Facility, EPFL, Switzerland) for help with crystallization, biochemical characterization, and providing SpCas9 protein. We thank Aline Aebi and Jean Philippe Gaudry (Bertarelli Foundation Gene Therapy Platform, EPFL, Switzerland) for their assistance with AAV production and purification. We thank Didier Nurizzo and Max Nanao (European Synchrotron Radiation Facility, MASSIF-3 and ID30B beamline, Grenoble, France) for assistance with crystallographic data collection. We thank Alexander Myasnikov, Bertrand Beckert, and Sergey Nazarov (Dubochet Center for Imaging, EPFL-UNIL-UNIGE, Switzerland) for assistance with cryo-EM data collection. We thank Miguel Garcia and Francesco Palumbo (Flow Cytometry Core Facility, EPFL, Switzerland) for their help with flow cytometry measurements. We thank the group of Ricardo Fernandes for generously providing purified CD45 ECD protein.

## Author contributions

M.P., L.N., C.S. and B.E.C. conceived the study and designed experiments. M.P., L.N., Y.C., C.A.G., and S.O. developed the code base. M.P., L.N. and C.S. generated protein designs. M.P., L.N., C.S., J.S., and S.G. purified proteins. L.N. and K.H.G. performed protein binding assays. E.P. and S.B. performed CD45 binder characterisation. K.H.G. and L.V. developed and performed PD-1 binder characterisation assays. G.N.H. purified SAS-6. L.K. performed gene editing assays. A.A-S. performed blocking assays for birch allergen. M.P. and L.N. solved crystal and cryoEM structures. P.B., A.H.W., and S.L. performed and analysed CbAgo activity assays and SEC analysis. S.K., J.C. and A.J.V. performed and analyzed Claudin experiments. C.S., L.N., B.S. and B.E.C designed and performed AAV experiments. B.J.Y., A.M.W., P.G., Y.D.M, G.S., D.C.S., A.J.V., S.O., B.S. and B.E.C. supervised the work and acquired core funding. M.P., L.N., C.S. and B.E.C. wrote the initial manuscript. All authors read and contributed to the manuscript. M.P., L.N. and C.S. agree to rearrange the order of their respective names according to their individual interests.

## Materials and Methods

### BindCraft design protocol

The input and design settings for running the BindCraft pipeline are organized into user-friendly JSON files. To initiate design trajectories, a target PDB format structure needs to be specified, along with the desired minimum and maximum length of the binders, and the desired number of final filtered designs. A target hotspot can be specified as either individual residues or entire chains, or can be omitted completely in which case a binding site is selected according to the combined design loss.

The binder hallucination process is performed using the ColabDesign implementation of AF2. The design process is initialized with a random sequence for the binder, which is predicted in single sequence mode, and a structural input template for the target. This is passed through the AF2 network to obtain a structure prediction and calculate the design loss. The design loss function is composed of multiple terms, with default weight values indicated in parentheses:

a. Binder confidence pLDDT (weight 0.1)
b. Interface confidence i_pTM (weight 0.05)
c. Normalized predicted alignment error (pAE) within the binder (weight 0.4)
d. Normalized predicted alignment error (pAE) between binder and target (weight 0.1)
e. Residue contact loss within binder (weight 1.0)
f. Residue contact loss between the target and binder - if hotspots are specified, the rest of the target is masked from this loss (weight 1.0)
g. Radius of gyration of binder (weight 0.3)
h. “Helicity loss” - penalize or promote backbone contacts every in a 3 residue offset to promote the hallucination of helical or non-helical designs (weight -0.3)
i. Optional “N&C termini loss” - increases the proximity of the N- and C-termini of the binder to allow splicing into protein loops (weight 0.1)

The loss function is used to calculate position specific errors, which are then backpropagated through the AF2 network to produce a L x 20 error gradient, where L is the sequence length. Using multiple iterations and stochastic gradient descent optimization, this error gradient is recomputed and used to optimize the input binder sequence for the next iteration to minimize the resulting loss. We backpropagate through the AF2 multimer model weights^19^ and swap randomly between the 5 trained models at each iteration to ensure robust sequence generation and reduce the risk of overfitting to a single model.

Since our goal is to arrive at a real discrete sequence for the binding interface, the sequence optimization is performed in four stages. The first sequence optimization stage is performed in a continuous sequence space using logit inputs. At each step, the sequence representation is based on linear combination of (1-λ) * logits + λ * softmax(logits/temp), where λ = (step+1)/iterations and temp = 1.0. Here, multiple amino acids are considered per each binder position, which allows the exploration of a larger and less constrained sequence-structure space. After 50 iterations, we terminate trajectories exhibiting poor AF2 confidence scores, as we found that such trajectories rarely converge to high confidence designs. Additionally, if a beta-sheeted trajectory is detected, we increase the number of recycles during design from 1 to 3 to ensure accurate prediction. The continuous sequence space optimization is then continued for additional 25 iterations. During the second optimization stage, the sequence logits are normalized to sequence probabilities using the softmax function for 45 iterations to funnel the design space towards a more realistic sequence representation defined as softmax(logits/temp) At each step, the temperature is lowered, where temp = (1e-2 + (1 - 1e-2) * (1 - (step + 1) / iterations)**2). The temperature is also used to scale the learning rate for rate decay. For the third stage, we implement the straight-through estimator, allowing the model to see the one-hot representation, but backprop through the softmax representation. This procedure is performed for 5 iterations. For the final fourth stage the sequence inputs are converted to a one-hot discrete encoding. At each step, X random mutations are independently sampled and tested from the probability distribution of the softmax representation from the previous stage, and mutations with best loss are fixed. X is defined based on the length of the binder sequence (0.05 * binder length). This procedure is performed for 15 iterations. At the end, trajectories with pLDDT below 0.7, less than 7 interface contacts, or significant backbone clashes are rejected.

Successful binder design trajectories are subjected to MPNN_sol_ sequence optimization to improve stability and solubility^20^. To this end, we preserve binder residues in a 4 Å radius around the target interface, and design 20 new sequences for the remaining binder core and surface residues using the soluble weights of ProteinMPNN^13^, with a temperature of 0.1 and 0.0 backbone noise. These optimized sequences are then re-predicted using the AF2 monomer model, with 3 recycles and 2-template based models^60^ in single sequence mode, to ensure robust and unbiased complex assessment. Each of the two resulting models is then energy minimized using Rosetta’s FastRelax protocol^61^ with 200 iterations, and interface scores are computed using the InterfaceAnalyzer mover^62^ with sidechain and backbone movement enabled.

Designs are finally filtered using a set of predefined filters to ensure the selection of high quality designs for experimental testing. Filters were initially defined based on experimental observations from previous binder design studies^7,8,10,11^ and refined over the course of this work. These include:

a. AF2 confidence pLDDT score of the predicted complex (> 0.8)
b. AF2 interface predicted confidence score (i_pTM) (> 0.5)
c. AF2 interface predicted alignment error (i_pAE) (< 0.35)
d. Rosetta interface shape complementarity (> 0.55)
e. Number of unsaturated hydrogen bonds at the interface (< 3)
f. Hydrophobicity of binder surface (< 35%)
g. RMSD of binder predicted in bound and unbound form (< 3.5 Å)

We allow only 2 MPNN_sol_ generated sequences per individual AF2 trajectory to pass filters to promote interface diversity amongst selected binders. This design procedure is set up to loop until a defined number of final desired designs is reached. For optimal results, we recommend running the design pipeline until at least 100 designs pass computational filters. This generally requires the sampling of about 300-3000 trajectories. We then usually pick 10 designs from the top 20 (ranked by i_pTM) for experimental testing.

### Design settings for individual target proteins

To generate designs against targets described in the results section, we utilized the following input structures, binder specifications, and hotspot designations. For AF2 predictions, we used full length input sequences from Uniprot. In all cases, the amino acid cysteine was excluded from sequence design. For AAV targets, the N&C termini loss is activated with default weight.

| *Target* | *Input PDB* | *Hotspots* | *Binder length* |
| --- | --- | --- | --- |
| PD-1 | AF2 prediction, trimmed to 32-146 | 64, 126, 129, 133 | 80-150 |
| PD-L1 | AF2 prediction, trimmed to 18-132 | 54, 56, 66, 115 | 65-155 |
| IFNAR2 | 2LAG<sup>63</sup>, trimmed to 8-110 | 52, 80, 82, 84, 96, 98 | 60-175 |
| CD45 | AF2 prediction of 5FMV<sup>64</sup>, split by extracellular fibronectin domains | d1, d2, d3, d4, d1-d2, d2-d3, d3-d4 | 80-200 |
| Claudin1 | AF2 prediction of CLN1-14<sup>20</sup> | 31, 46, 55, 152 | 80-175 |
| BBF-14 | 9HAG | none | 70-250 |
| CrSAS-6 | AF2 prediction, trimmed to 15-160 | none | 90-200 |
| Der f7 | AF2 prediction with 3UV1 template<sup>65</sup>, trimmed to 18-213 | 132 or 178 or 201 | 70-185 |
| Der f21 | AF2 dimer prediction with 5YNY template<sup>32</sup>, trimmed to 25-136 | 34 or 52 or 87 or 105 or 116 | 70-185 |
| Bet v1 | AF2 prediction | 24 | 70-185 |
| SpCas9 | AF2 prediction with 4ZT0<sup>39</sup> template, trimmed to 96-174 + 306-446 | 360 | 70-150 |
| CbAgo | AF2 prediction with 6QZK template<sup>46</sup>, trimmed to 1-175 + 298-343 + 635-689 (N+PIWI) or 180-276 (PAZ) | 39 (N+PIWI) or 268 (PAZ) | 70-160 |
| HER2<br>(AAV) | 1N8Z<sup>66</sup>, trimmed to 23-268 | 170 or 170, 165 | 60-100 |
| PDL1<br>(AAV) | AF2 prediction, trimmed to 18-132 | 54, 56, 58, 63, 66,<br>113, 115 | 75-115 |

### Computational benchmarks of BindCraft

To evaluate the flexibility of the target structure post-design, the input PDB structure of the target was aligned to the target chain A of the design trajectory, and RMSD_Cα_ was calculated using PyRosetta. For increasing target flexibility, the sequence of the input target template was masked by enabling the flag ‘rm_target_seq’ in ColabDesign for trajectory hallucination^67^, and 200 trajectories were generated. For the impact of the helicity loss on binder secondary structure composition, the ‘weights_helicity’ flag in BindCraft was set to 1, 0, -0.3, -1, -2, and -3, and 200 trajectories were generated for each instance using otherwise default settings. To compare the design capabilities of AF2 monomer and multimer weights, we generated 200 trajectories each. For AF2 multimer trajectories, we used the default settings where AF2 multimer models 1-5 are used for design and AF2 monomer models 1-2 trained with templates are used for reprediction. For AF2 monomer this is inverted, we use AF2 monomer models 1 and 2 for design and AF2 multimer models 1-5 for reprediction. For benchmarks involving design and trajectory success rates, we run the design pipeline either for 200 trajectories or until 100 designs passing *in silico* filters are accumulated (where indicated). We then designate trajectories with pLDDT above 0.7 as ‘Passing’, while trajectories that have pLDDT below 0.7, more than one Cα backbone clash between chains or less than 3 contacts between the binder and target as ‘Low Confidence’. Benchmarking of designs from other design pipelines was performed using the BindCraft prediction method of either AF2 monomer or multimer in single sequence, with templates provided for the target according to the specifications in their respective publications.

### Protein expression, purification, and characterization

DNA sequences of designed proteins, as well as BBF-14, Der f7, Der f21, and Bet v1 targets were ordered from Twist Biosciences with Gibson cloning adapters for cloning into bacterial expression vectors pET21b or pET11. Proteins were expressed in *Escherichia coli* BL21 Codon Plus (DE3) cells (Novagen) by inducing with 0.5 mM IPTG for 6 hours at 18 °C. Pellets were resuspended and lysed in lysis buffer (50 mM Tris-HCl pH 7.5, 500 mM NaCl, 5% glycerol, 1 mg/ml lysozyme, 1 mg/ml PMSF and 1 µg/ml DNAse) using sonication. Cell lysates were clarified using ultracentrifugation, loaded on a 1 ml Ni-NTA Superflow column (QIAGEN) and washed with 7 column volumes of 50 mM Tris-HCl pH 7.5, 500 mM NaCl, 10 mM imidazole. Proteins were eluted with 10 column volumes of 50 mM Tris-HCl pH 7.5, 500 mM NaCl, 500 mM imidazole. Claudin binders were dialyzed against 20 mM HEPES pH 8.0, 150 mM NaCl, 4% glycerol and directly frozen.

Fc-fused PD-L1 target^8^, IFNAR2 target, the IFNA2 cytokine, and antibodies were expressed using a mammalian Expi293 secreted expression system (Thermo Fisher Scientific, A14635). Six days post transfection the supernatants are collected, cleared and purified either using a 1 ml Ni-NTA Superflow column (QIAGEN) or protein A affinity column (QIAGEN). SAS-6^27^, SpCas9^68^, CbAgo and the catalytic mutant of CbAgo (D541A, D611A)^46^ have been purified as described previously.

Remaining bacterial and mammalian expressed proteins were then concentrated and injected onto a Superdex 75 16/600 gel filtration column (GE Healthcare) in 50 mM Tris-HCl pH 7.5, 250 mM KCl. Proteins after size exclusion were concentrated, frozen in liquid nitrogen, and stored at -80 °C. Molar mass, sample homogeneity, and multimeric state were confirmed using SEC-MALS (miniDAWN TREOS, Wyatt) by injecting 100 µg of protein in PBS. Folding, secondary structure content, and melting temperatures were assessed using circular dichroism in a Chirascan V100 instrument from Applied Photophysics in PBS at a concentration of 0.1-0.3 mg/ml.

### Expression and purification of PD-1 target and binders

DNA sequences were synthesized in the pcDNA3.4 vector with an osteonectin secretion signal at the N-terminus (Twist Biosciences). *De novo* designs were fused to the N-terminus of human IgG1 Fc. The extracellular domain (25-167) of human PD-1 (UniProtKB: Q15116) was fused to a C-terminal AviTag™ and His tag. Plasmid DNA was prepared from glycerol stocks (Twist Biosciences) using Cowin Biosciences GoldVac EndoFree plasmid maxi kit. Plasmids were transfected into 3 mL or 50 mL cultures of Expi293F™ (Gibco) cells per the manufacturer’s recommendations. Cells incubated at 37 °C for 4-5 days prior to harvest. Following protein expression, the cell culture supernatant was filtered through a 0.22 µM filter and purified using MabSelect protein A affinity chromatography resin (Cytiva). The column was washed with PBS and the protein was eluted in Tris glycine buffer pH 2.5. Following elution, proteins were dialyzed into PBS using a 10 kDa MWCO dialysis cassette. For production of biotinylated PD-1 protein, the PD-1 plasmid was co-transfected with BirA plasmid (2:1 ratio). The BirA plasmid contains the BirA sequence (UniProtKB: P06709) with a C-terminal Flag tag in the pcDNA3.4 vector.

### Binding characterization of PD-1

Designs were initially screened for binding to biotinylated human PD-1 or a random protein using biolayer interferometry (Sartorius OctetRED384). Biotinylated human PD-1 protein and biotinylated lysozyme (GeneTex) were prepared at 500 nM in PBS containing 0.1% BSA (PBSA). The designs were diluted to 5 µM in PBSA. Streptavidin-labeled biosensors were saturated with either biotinylated human PD-1 or biotinylated chicken lysozyme. The designs were then allowed to associate with the immobilized ligand for 60 seconds, followed by a dissociation step in PBSA. The baseline subtracted signal (nm) was calculated and used to prioritize human PD-1 specific binders for further characterization.

To determine the affinity of selected designs, 100 nM biotinylated human PD-1 prepared in PBSA was immobilized onto a streptavidin labeled biosensor for 15 seconds. Serial dilutions of the designs (from 2.5 µM to 5 nM) were then allowed to associate with the immobilized ligand for 180 seconds, followed by a dissociation step in PBSA for 300 seconds. Following background subtraction of the BLI binding curves using the buffer only (PBSA) curve, the K_d_ was determined using the 1:1 model in the Data Analysis HT 11.1 curve fitting module.

To determine if the designed protein competed with pembrolizumab for binding to PD-1, 100 nM biotinylated human PD-1 in PBSA was immobilized onto streptavidin coated biosensors for 15 seconds. An initial association with 200 nM pembrolizumab prepared in PBSA was performed for 180 seconds, followed by a second association with 200 nM design prepared in PBSA for 180 seconds.

### Surface Plasmon Resonance (SPR) binding and competition assays

SPR measurements were performed using the Biacore 8K system (Cytiva) in HBS-EP+ buffer (10 mM HEPES pH 7.4, 150 mM NaCl, 3 mM EDTA, 0.005% (v/v) Surfactant P20 GE Healthcare). Target proteins were immobilized on a CM5 chip (GE Healthcare) through amide coupling in 10 mM NaOAc pH 4.5 for 250s at a flow rate of 10 µl/min aiming for 100 relative response units. Designed binders or control proteins were injected as analytes in either a single 10 µM concentration during binder pre-screening or in serial dilutions to assess binding kinetics. These were injected at a flow rate of 30 µl/min for a varying contact time, followed by dissociation. If necessary, the chip surface was regenerated after each injection using 10 mM Glycine-HCl pH 2.5 for 30s at a flow rate of 30 µl/min. Binding curves were fitted with a 1:1 Langmuir binding model in the Biacore 8K analysis software. Steady-state response units were plotted against analyte concentration and a sigmoid function was fitted to the experimental data in Python 3.9 to derive the K_D_.

Competition assays were performed as follows. For PD-L1 and IFNAR2: Target receptors were immobilized, and binders and competitors were injected as analytes. Two subsequent injections were performed either with only competitor (A,1 µM), only design (B,1 µM) or first competitor (1 µM, A) and then design+competitor (both1 µM, A+B). For BetV1: REGN5713 (Antibody format) was immobilized on the SPR chip and in a first injection (1) loaded with BetV1 allergen (1 µM), before either REGN5714 (Fab format) or Birch_binder2 were injected (both 1 µM) (2).

### Protein crystallization and structure determination

The BBF-14_binder4 complex was crystallized at a concentration of 5 mg/ml using sitting drop vapor diffusion at 16 °C in 0.1 M MES pH 6.0, 0.2 M Na acetate trihydrate, 20% w/v PEG 8000 buffer (SG1-Eco Screen, Molecular Dimensions). The DerF7_binder2 complex in P21 crystal form was crystallized at a concentration of 15 mg/ml using sitting drop vapor diffusion at 16 °C in 0.1 M MES pH 6.5, 0.2 M KSCN, 25% w/v PEG 2000 MME buffer (Clear Strategy Screen I, Molecular Dimensions). The DerF7_binder2 complex in C121 crystal form was crystallized at a concentration of 15 mg/ml using sitting drop vapor diffusion at 16 °C in 0.1 M MES pH 6.5 and 20% v/v PEG smear high BCS (BCS Screen, Molecular Dimensions). The DerF21_binder10 complex was crystallized at a concentration of 30 mg/ml using sitting drop vapor diffusion at 16 °C in 0.1 M Na citrate pH 5.6, 1.0 M LiSO_4_, 0.5 M NH_4_SO_4_ buffer (SG1-Eco Screen, Molecular Dimensions). Crystals were cryoprotected in 25% glycerol and flash-cooled in liquid nitrogen. Diffraction data was collected at the European Synchrotron Radiation Facility MASSIF-3 and ID30B beamlines, Grenoble, France at a temperature of 100 K. Crystallographic data was processed using the autoPROC package^69^. Phases were obtained by molecular replacement using Phaser^70^. Atomic model refinement was completed using COOT^71^ and Phenix.refine^70^. The quality of refined models was assessed using MolProbity^72^. Structural figures were generated using ChimeraX^73^.

### CryoEM structure determination

SpCas9 was mixed with a 3-fold excess of either binder3 or binder10, and the complex was purified using S200 10/300 gel filtration column (GE Healthcare) in 20 mM Tris-HC pH 7.5, 250 mM KCl. The purified complex was applied to a glow discharged 300-mesh holey carbon grid 300-mesh holey carbon grid (Au 1.2/1.3 QuantifoilMicro Tools), blotted for 4 seconds at 95% humidity, 10 °C, plunge frozen in liquid ethane (Vitrobot Mark IV, FEI) and stored in liquid nitrogen. Data collection was performed on a 300 kV Titan Krios G4 microscope equipped with a FEI Falcon IV detector and SelectrisX energy filter. Micrographs were recorded at a magnification of 165kx, pixel size of 0.726 Å, and a nominal defocus ranging from -0.8 mm to -2.2 mm.

Acquired cryo-EM data was processed using cryoSPARC v4.5.3^74^. Micrographs were patch motion corrected, and micrographs with a resolution estimation worse than 5 Å were discarded after patch CTF estimation. Initial particles were picked using blob picker with 90-135 Å. Particles were extracted with a box size of 360×360 pixels, down-sampled to 220×220 pixels. After 2D classification, clean particles were used for *ab initio* 3D reconstruction and initial non-uniform 3D reconstruction^75^. This model was used for additional template-based picking of particles. Following several rounds of 3D classification, where classes containing unbound Cas9 were excluded, the class with most detailed binder features was re-extracted using full box size and subjected to non-uniform and local refinement to generate final reconstructions. The local resolution was calculated and visualized using ChimeraX^73^. The *in silico* models were docked into density using ChimeraX^73^.

### Birch allergen blocking assay

Anti-Bet v1 binder blocking capacity was assessed by first coating NuncSorp (Thermofisher) plates with 2 μg/ml of anti-human IgE monoclonal antibody (NBS-C BioScience, Vienna, Austria; clone Le27; Cat#0908-1-010) in coating buffer (15 mM Na_2_CO_3_, 34.87 mM NaHCO_3_) and incubating overnight at 4 °C. The plates were washed with PBS+0.05% Tween and blocked using PBS+1% BSA for 2 hours at room temperature. Then, sera of birch allergic patients were added at a concentration of 4 ng/ml of anti-Betv1 IgE. Biotinylated Bet v1 allergen at 1 nM concentration was preincubated for 2 hours at room temperature with 4-fold serial dilutions of the BetV1_binder2 starting at 2 μM or with 5-fold serial dilutions of the cocktail of REGN5713, REGN5714 and REGN5715 (starting at 50 nM each) and then added to the IgE coated plate. After two-hour incubation at room temperature, the plates were washed with PBS+0.05% Tween and streptavidin horseradish peroxidase (BD Pharmigen, USA; Cat#554066; 1:1000 dilution) was added and incubated for 1 hour. Plates were washed and tetramethylbenzidine substrate (BD Biosciences, San Diego, USA; Cat#555214) was added and incubated further 20 minutes. The reaction was stopped with 2 M sulfuric acid. Absorbance was measured on a spectrophotometer at 450 nm with a 630 nm reference, and blocking percentage was measured by subtracting the absorbance of the sample in the absence of the binder.

### Cy5 labelling of claudins

CLDN1 WT was labelled with Cy5 by adding a 1:5 molar excess of dye and incubating for 2 h on ice. The excess dye was removed by passing through a PD-10 column. The labelled protein was collected and stored in small aliquots at -80 C after flash freezing in liquid nitrogen.

### Microscale Thermophoresis (MST)

For MST based interaction studies, the Monolith (Nanotemper) instrument was used. Serial dilutions of the ligand (CLDN1_b12/CpE_Nd33) were made in buffer B (25 mM HEPES pH 8.0, 200 mM NaCl, 5% glycerol 0.03% DDM) and mixed with 10 nM of labelled CLDN1 WT. After 10 min of incubation, samples were transferred to capillaries (Monolith standard capillary) and readings were initiated. The spectral shift data was plotted and fitted into a Kd model and estimated Kds were obtained. When data was not fitted using the K_d_ model, Hill model was used to fit data. For studying the competitive binding of CpE_Nd33 and CLDN1_b12 to the target CLDN1 WT, a second set of experiments was performed. CLDN1 WT was incubated with CLDN1_b12 (2X of K_d_) and subsequently challenged with CpE_Nd33.

### Cytotoxicity assay

To study if claudin bidners were able to inhibit pore-formation in Sf9 cells expressing claudins, adherent Sf9 cells in a 24-well plate were infected with baculovirus containing either CLDN1 or CLDN4. The assay was performed as shown previously^25^. Briefly, for each claudin, a 12-well experiment was performed. Six wells were used to test the effect of binders on the pore-forming capacity of CpE_Nd33 and the other six wells were used as controls. After 36 h of infection, 4 µM of each binder were added into six different wells and the plate was then gently mixed by swirling and incubated for 5 min. After that, 300 nM of CpE_Nd33 was added to each of the six wells. The following controls were used in experiment 1. Sf9 without baculovirus infection, 2. Sf9 infected with claudin but not treated with CpE_Nd33, 3. Sf9 infected with Claudin and treated with CpE_Nd33 4. Sf9 infected with Claudin and treated with COP4 Fab (referred to as CpE Inhibitor) 5. Sf9 infected with Claudin and treated by COP4 followed by addition of CpE_Nd33 after incubation for 5 min. The number of cells dead or alive were then measured after 18 h of incubation by staining the cells with trypan blue and measuring the number of cells using an automated cell counter (Invitrogen Countess).

### SpCas9 gene editing

For SpCas9-sgRNA plasmid cloning, lentiCRISPR v2 (Addgene #52961, a gift from Feng Zhang) was digested with BsmBI (NEB). Oligonucleotides encoding for the sgRNA targeting the NSD2 gene were annealed and ligated into the digested lentiCRISPR v2 plasmid. All binders were human codon optimized using the GenSmart Codon Optimization tool and ordered as inserts with homology overhangs for cloning from Twist bioscience. Final binder plasmids were generated by isothermal assembly (NEBuilder® HiFi DNA Assembly Cloning Kit, NEB).

HEK293T (ATCC CRL-3216) were maintained in DMEM plus GlutaMax (Thermo Fisher Scientific), supplemented with 10% (vol/vol) fetal bovine serum (FBS, Sigma-Aldrich) and 1 × penicillin-streptomycin (Thermo Fisher Scientific) at 37 °C and 5% CO2. Cells were maintained at confluency below 90% and passaged every 2-3 days. For testing inhibitor efficiency, HEK293T cells were seeded in 48-well cell culture plates (Greiner) and transfected at 70% confluency using 300 ng Cas9+sgRNA plasmid, 500 ng inhibitor plasmid, and 5 uL Lipofectamine 2000 according to the manufacturer’s instructions (Thermo Fisher Scientific). The next day, cells were split and selected with either Puromycin, Blasticidin or both. Three days post-transfection, cells were harvested and genomic DNA was isolated by direct lysis.

The DNA from the cell lysate was prepped for next-generation sequencing as previously described^76^. In the first PCR round, genomic regions of interest were amplified using GoTaq Green Master Mix (Promega) and primers that included Illumina forward and reverse adaptor sequences. A second PCR round, also using GoTaq Green Master Mix (Promega), introduced p5-p7 barcodes into the products from the first round. The resulting amplified amplicons were pooled and quantified using a Qubit 3.0 fluorometer (Invitrogen). The libraries were then sequenced using a MiSeq platform (Illumina, 150 bp, paired-end). Sequencing data and resulting gene editing insertion-deletion rates were analyzed using CRISPResso2^77^.

### CbAgo in vitro cleavage assay

For *in vitro* cleavage assays, binders, CbAgo, 5’-phosphorylated 16 nt ssDNA guide (oDS423), and Cy5-labelled 45 nt ssDNA target (oDS401) were mixed to final concentrations of 2:0.4:0.4:0.2 μM in 10 mM HEPES pH 7.5, 125 mM KCl, and 2 mM MgCl_2_. To this end, first the binder protein and CbAgo were mixed and incubated at 37°C for 15 minutes, after which the mixture was incubated on ice and guide ssDNA and target ssDNA were added. Subsequently, reaction mixtures were incubated at 37 °C, and samples were taken at 0, 4, 10, 30, and 60 minute timepoints. Samples taken at each timepoint were directly quenched by adding 2x RNA loading dye (25 mM EDTA, 5% v/v glycerol, 90% v/v formamide) and heating for 5 minutes at 95 °C. Cleavage products were resolved using denaturing (7M Urea) 20% polyacrylamide gel electrophoresis, and gels were imaged using a Amersham Typhoon gel scanner (Cytiva Life Sciences). Cleavage reactions were performed in triplicates for each binder protein.

CbAgo target cleavage was quantified using ImageQuant TL 1D v8.2.0 (Cytiva Life Sciences), and fitted with non-linear least squares fit (nlsLM from R package minpack.lm) to a double-exponential decay model to model initial (fast) and turnover (slow) cleavage:

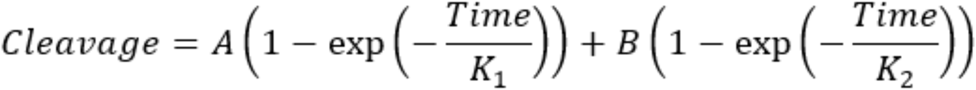

If fitting to a double-exponential decay model yielded no fit after 1024 iterations with residuals and gradient convergence tolerance of 1*10^-9^, the turnover cleavage (slow) was considered negligible, and a single-exponential decay model (i.e. B=0) was used.

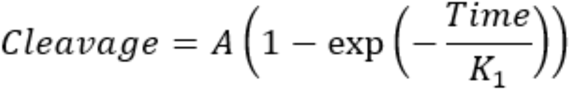

For all samples K_cat_ was calculated from the fit constants for the initial rate (A and K_1_):

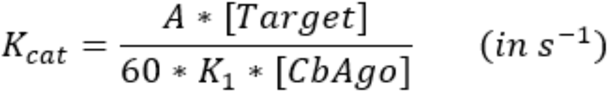

The mean and standard deviation of K_cat_ was calculated using the three experimental replicates.

### CbAgo biolayer interferometry binding kinetics

Biolayer Interferometry measurements were conducted using the Gator BLI system and GatorOne software (Gator Bio, version 2.7.3.0728). The running buffer consisted of 150 mM KCl, 20 mM HEPES (pH 7.5), and 0.5% BSA. His-tagged CbAgo binders were immobilized on the sensor tips at a concentration of 10 µg/mL. After immobilization, the tips were transferred into serial dilutions of CbAgo. Binding curves were globally fitted using a 1:1 interaction model in the Gator software.

### CbAgo size exclusion chromatography (SEC) binding verification

Purified CbAgo was diluted to 0.8 mg/mL (9.3 µM) and mixed with 0.2 mg/mL binder protein and 9.3 µM 5’-phosphorylated 16 nt ssDNA guide (oDS423) in SEC buffer (20 mM HEPES pH 7.5, 250 mM KCl, and 2 mM MgCl_2_). The mixture was incubated for 15 minutes at room temperature. After incubation samples were resolved at RT on a Superdex 200 Increase 10/300 GL column (Cytiva Life Sciences) connected to a 1260 Infinity II HPLC system (Agilent) using SEC buffer with a flow-rate of 0.75 mL/min. The elution was measured using a Agilent 1260 Infinity II Multiple Wavelength Detector at 280 nm. The data were analysed using Astra 8.1 (Wyatt Technology).

### AAV Engineering

HEK293 cells adapted to culture in orbitally shaken bioreactors (HEKExpress™, ExcellGene SA) were maintained in Serum-free BalanCD® HEK293 Medium (Irvine Scientific) supplemented with L-alanyl-L-glutamine (Gibco Glutamax) at 37°C, 80% humidity, 5% CO_2_, under constant shaking at 180 rpm (shaking diameter 5 cm). Cells were passaged every 3-4 days to a concentration of 0.2*10^6^ cells/mL. For the generation of cell lines stably expressing the target receptors used in the AAV transduction experiments, the receptor cDNAs were obtained from an ORF collection (HER2: ORFeome Collaboration (OC) cDNA Clone, PDL1: Addgene #121142) and cloned into a pRRLSIN lentiviral shuttle construct (Addgene #12252) with expression under the control of the human phosphoglycerate kinase (hPGK) promoter. Lentiviral particles were generated using standard procedures for calcium phosphate transfection of HEK293T cells with the pRRLSIN-hPGK-WPRE, p8.92, pMD2G and pAdVAntage™ plasmids. At 48 h, the vector-containing supernatant was harvested, filtered and concentrated by ultracentrifugation. The amount of lentiviral particles present in the obtained vector suspension was quantified using a p24 antigen ELISA kit (ZeptoMetrix). HEKExpress™ cells were transduced in a 6-well plate at a density of 3.0*10^6^ cells/well using a multiplicity of infection (MOI) of 100 vg/cell (conversion factor: 1 pg p24 = 1e4 vg). After 5 days, the cells were stained for the presence of the respective target receptor using an APC-conjugated antibody (0.8 µg/mL, BioLegend, #329707 (PDL1), #324407 (HER2)) in staining buffer (PBS containing 0.5% BSA (Merck)) and sorted by flow cytometry using a Sony SH800 cell sorter. After expansion, the cells were aliquoted and frozen at -80 °C until further usage.

The pRepCap plasmids for the AAV production by transient transfection of HEKExpress™ cells, encoding the rep (AAV2) and cap (varying) genes, were chosen according to the different variants as indicated. For serotype 6 wild-type AAV (WT), an AAV6 plasmid was ordered from the manufacturer (Aldevron, pALD-AAV6). For the variant carrying the knock-out (KO) mutations to deplete the primary interactions with heparin (K459S and K531E) and sialic acid (V473D, N500E and T502S), a corresponding gene fragment was ordered as an insert with homology overhangs (Twist Biosciences) and cloned into pALD-AAV6 by BspEI/MscI yielding the pRepCap KO. To introduce the sequence encoding the designed miniprotein binders, an intermediate plasmid was created which, in addition to the five mutations to deplete the primary interactions, carries two silent mutations yielding MluI/NheI restriction sites in proximity to the chosen site of binder insertion between amino acid positions 497 and 498 of the serotype 6 VP3. The DNA sequences for the designed miniprotein binders, flanked by a single -(GSG)1-linker at both termini, were human codon optimized using the GenSmart Codon Optimization tool and ordered as inserts with homology overhangs (Twist Biosciences) for subcloning.

For the AAV production for screening in small-scale, the cells were seeded in 24-well cell culture plates at a density of 0.4*10^6^ cells/mL in a volume of 500 µL, and transfected with 520 ng pHelper (Aldevron, pALD-HELP), 250 ng shuttle plasmid (Aldevron, pALD-ITR-GFP), 270 ng pRepCap (varying) and 1.5 µg polyethyleneimine (PEI) (Polysciences). If applicable, the variants’ pRepCap plasmids were respectively mixed in a ratio of 1:2 with the pRepCap KO plasmid. 12 h after transfection, the cells’ media was exchanged and supplemented with 4 mM Valproic acid (VPA) (Sigma). The cell culture was incubated at the standard conditions described above but without shaking, and the AAV containing medium was harvested on day 5 by collecting the supernatant using centrifugation at 400 g for 5 min at room temperature to remove cells.

For the AAV production for validation at a normalized multiplicity of infection (MOI), the cells were seeded at a density of 1.0*10^6^ cells/mL in a volume of 300 mL in a TubeSpin 600 bioreactor tube (TPP) and transfected with 231 µg pHelper (Aldevron, pALD-HELP), 105 µg shuttle plasmid (Aldevron, pALD-ITR-GFP), 105 ng pRepCap (varying) and 900 µg polyethyleneimine (PEI) (Polysciences). If applicable, the pRepCap plasmids were obtained as described above, and mixed in a ratio of 1:2 with the pRepCap KO plasmid. 6 h after transfection, the cell culture medium was supplemented with 4 mM VPA (Sigma). The cell culture was incubated at the standard conditions described above for 7 days, and the vector was harvested on day 3-4 and on day 7 by collecting the supernatant after centrifugation of the bioreactor tube at 800 g for 10 min at room temperature. The supernatant was filtered (Stericup® Quick Release, Millipore Express® PLUS 0.22 μm PES, 1000 mL, Merck Millipore). The particles were concentrated from the cell culture supernatant to a concentration of at least 3.0*10^10^ vg/mL via Amicon® Ultra-15 centrifugal filter units, MWC 100 kDa (Merck). The WT and KO variant particles were processed alternatively according to Gaudry et al^78^. In short, the particles were purified from the cell culture supernatant using the POROS™ CaptureSelect™ AAVX resin (Thermo Fisher Scientific) on an ÄKTA Pure chromatography system followed by buffer exchange to PBS, 0.001% Pluronic® F-68 (10% stock solution, Gibco) via Amicon® Ultra-15 centrifugal filter units, MWC 100 kDa (Merck). The amount of genome-containing AAV particles was determined after treatment with DNase I (ThermoFisher) by digital PCR (dPCR) using the QIAcuity system and PCR kit (Qiagen).

For transduction, the target cells were seeded in 96-well cell culture plates at a density of 0.3*10^5^ cells/mL in a volume of 100 µL. After 6 h, the cells’ media was replaced with 100 µL AAV containing medium from the production in 24-well cell culture plates, or a 100 µL dilution to 3*10^10^ vg/mL of the material from the production in 300 mL culture to yield a MOI of 1*10^5^ vg/cell. If applicable, 0.8 µg/mL target receptor blocking antibody (BioLegend, #329707) was added. After 48 h, the cells were washed twice with 100 µL PBS containing 0.5% BSA (Merck), and the transduction signal (GFP) was measured by flow cytometry on an Attune NxT analyzer (ThermoFisher) equipped with an automated plate reader. The results were analyzed using FlowJo™ v10.8 Software (BD Life Sciences).

## Extended Data

**Extended Data Fig. 1.**
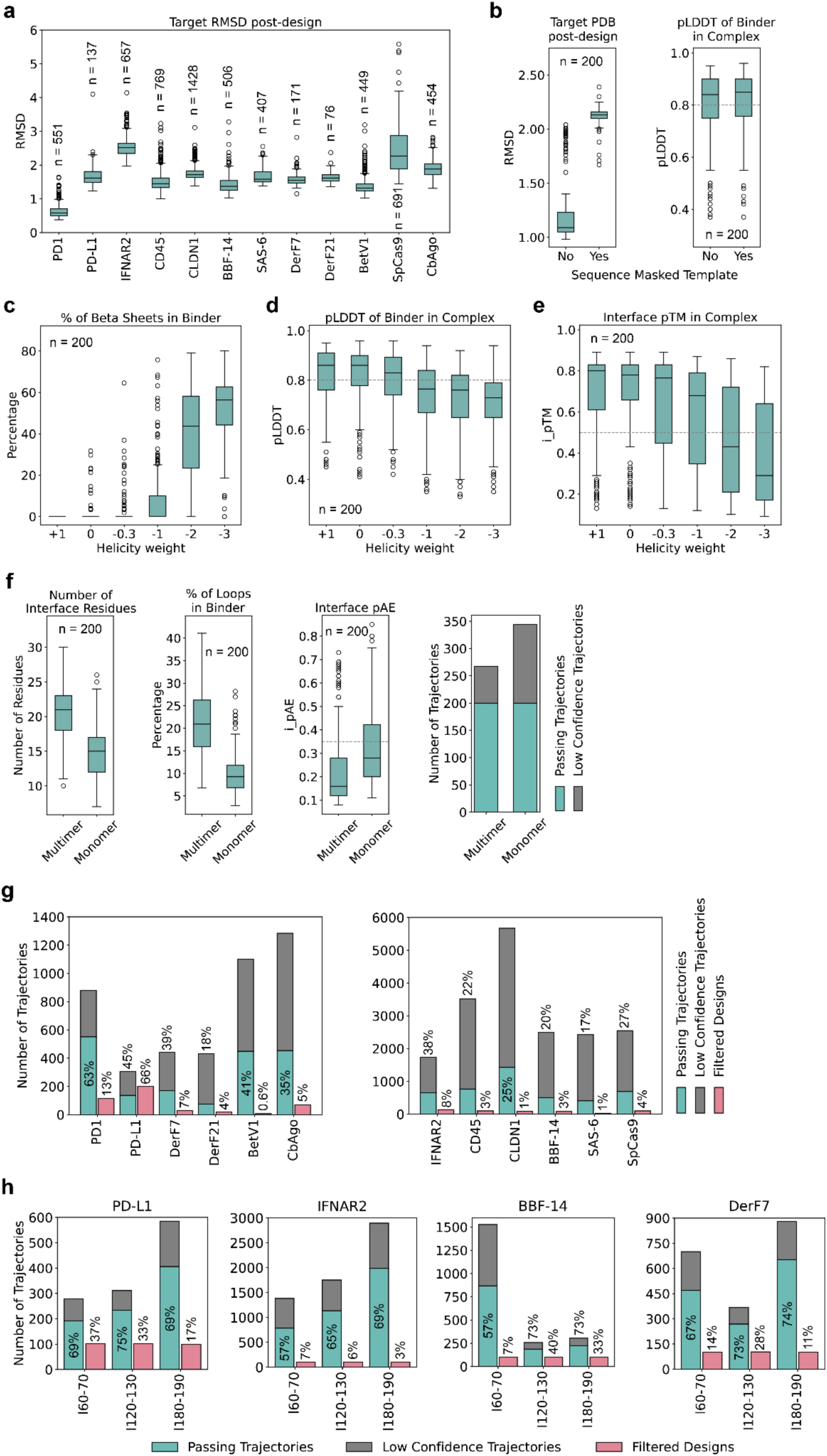
*In silico* benchmarking of BindCraft designs. **a,** Graphs plotting the RMSD_Cα_ of the target structure after initial AF2 binder hallucination, compared to the input target structure for different design targets. **b,** The RMSD_Cα_ and pLDDT confidence of the hallucination trajectory for PD-L1 with and without sequence masking of the input template. Sequence masking allows for higher levels of backbone flexibility. **c,** Effect of the helicity loss on secondary structure content of binders designed against PD-L1. Negative values that discourage the formation of alpha-helices can result in purely beta sheeted binders, however on average with reduced **d,** complex and **e,** interfaces confidences. **f,** Comparison of AF2 multimer- and monomer-based hallucination of binders against PD-L1. AF2 multimer interfaces are on average bigger, have a higher proportion of loops and higher interface confidence. **g,** Proportion of ‘Passing’ and ‘Low Confidence’ trajectories, alongside final designs passing all *in silico* filters for all design targets. Percentage above trajectory indicates proportion of Passing vs Low Confidence trajectories. Percentage above Filtered Designs indicates proportion of final passing designs vs total number of trajectories. **h,** Benchmark of design success rates depending on desired binder length across four targets. Benchmarks were run until 100 designs passed *in silico* filters.

**Extended Data Fig. 2.**
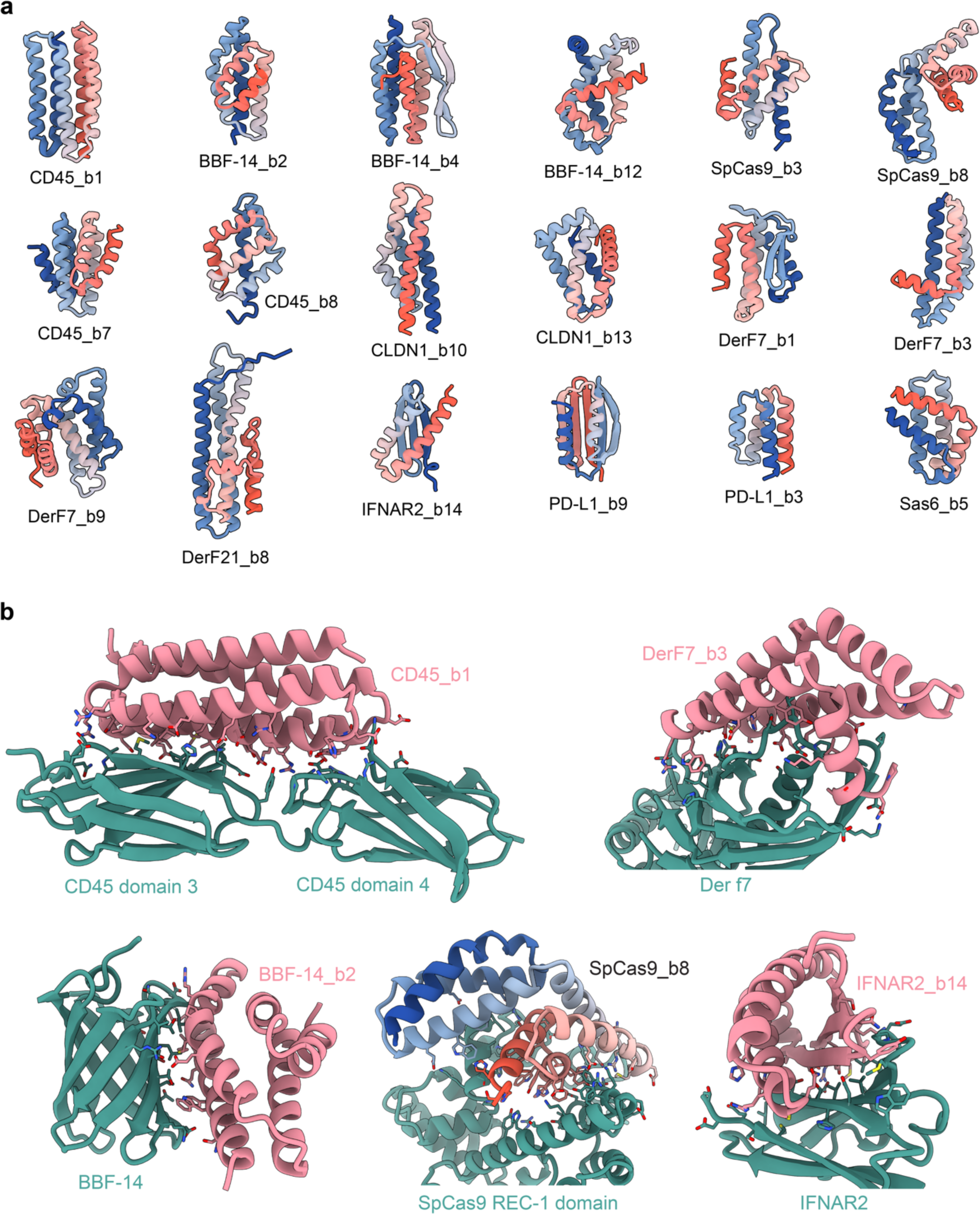
Examples of experimentally validated BindCraft designs. **a,** Cartoon representations of experimentally validated BindCraft binder topologies, coloured from N- (blue) to C-terminus (red). **b,** Examples of BindCraft generated interfaces between the binders (salmon or blue-red) and the target (green). Interface residues are highlighted.

**Extended Data Fig. 3.**
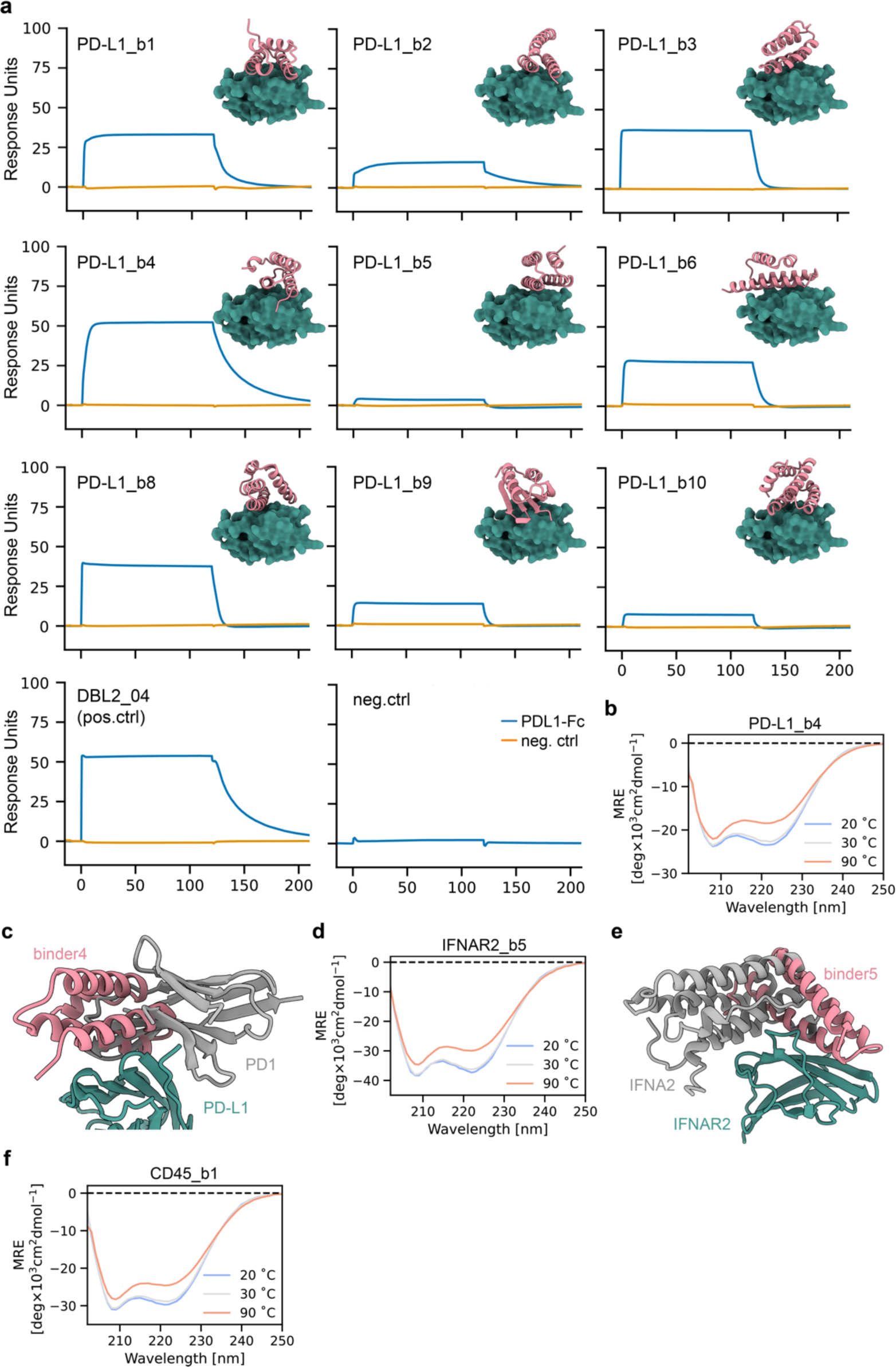
Biochemical characterization of binders targeting cellular receptors. **a,** SPR pre-screening of PD-L1 binder designs at 10 µM against immobilized PD-L1-Fc (blue) and a negative control (orange). The previously reported DBL2_04^8^ was included as a positive control. **b,** CD spectra of PD-L1 binder4 at different temperatures. **c,** Structural overlay highlighting the binding of mode of PD1 (gray) versus PD-L1_b4 (salmon). **d,** CD spectra of IFNAR2 binder5 at different temperatures. **e,** Structural overlay highlighting the binding of mode of IFNA2 (gray) versus IFNAR2_b5 (salmon). **d,** CD spectra of CD45 binder 1 at different temperatures.

**Extended Data Fig. 4.**
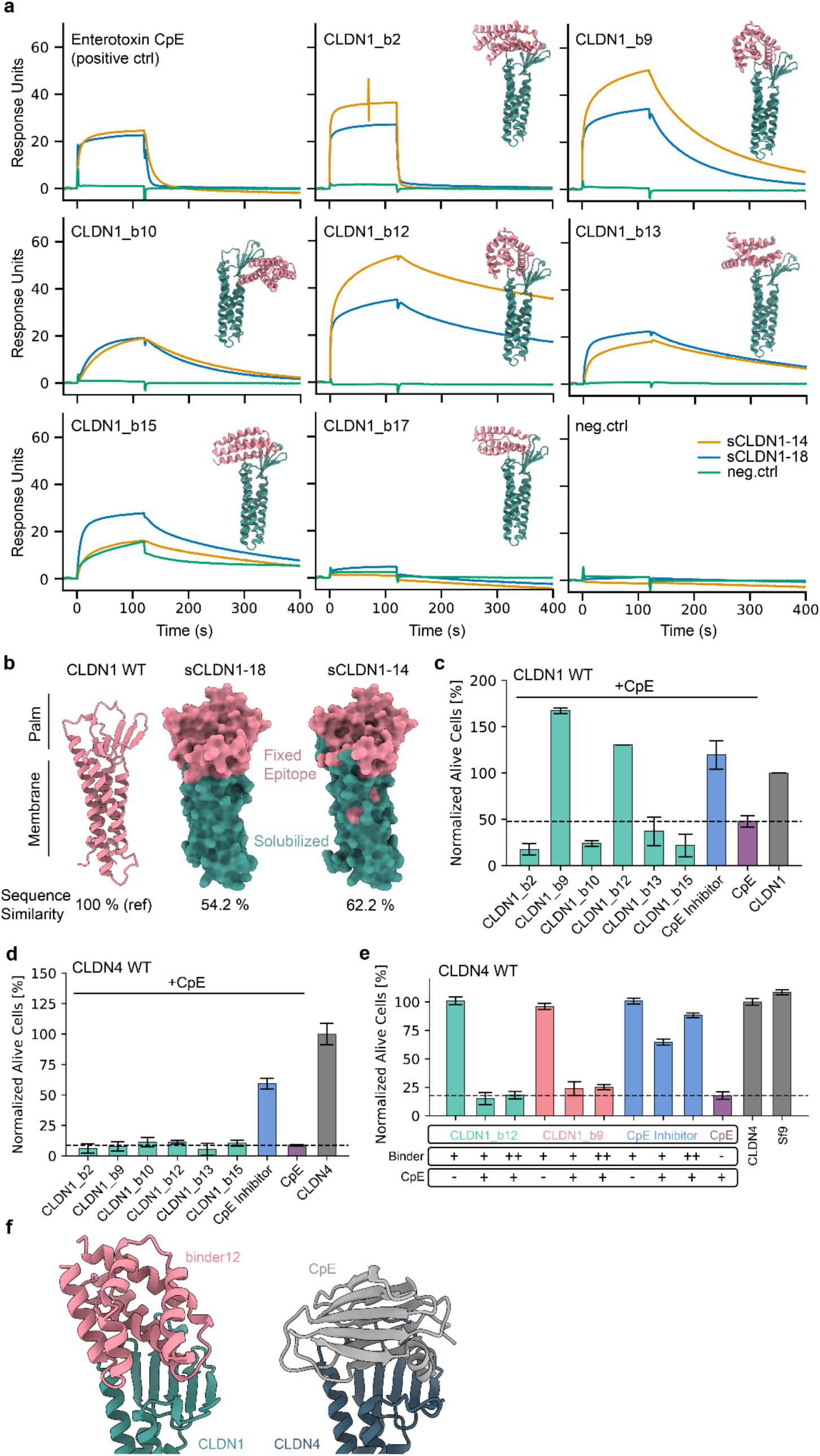
Biochemical characterization of Claudin binders. **a,** SPR pre-screening of CLDN1 binder designs at 10 µM against immobilized soluble analogues (sCLDN1-18:blue, sCLDN1-14:orange) and a negative control (green). The enterotoxin CpE was included as a positive control. **b,** Epitope comparison of CLDN1 and designed soluble analogues. Preserved wild type CLDN1 epitope residues on soluble analogues are indicated in pink. **c,** Cell-based screening assay for inhibition of CpE cytotoxicity in the context of CLDN 1. **d,** Cell-based assay showing concentration dependent inhibition of CpE cytotoxicity by a CpE Inhibitor, but not binder9 and binder12 in the context of CLDN4. **e,** Cell-based screening assay for inhibition of CpE cytotoxicity in the context of CLDN4. **f** Structural overview of the binding mode of binder12 (salmon) and the native CpE toxin (gray).

**Extended Data Fig. 5.**
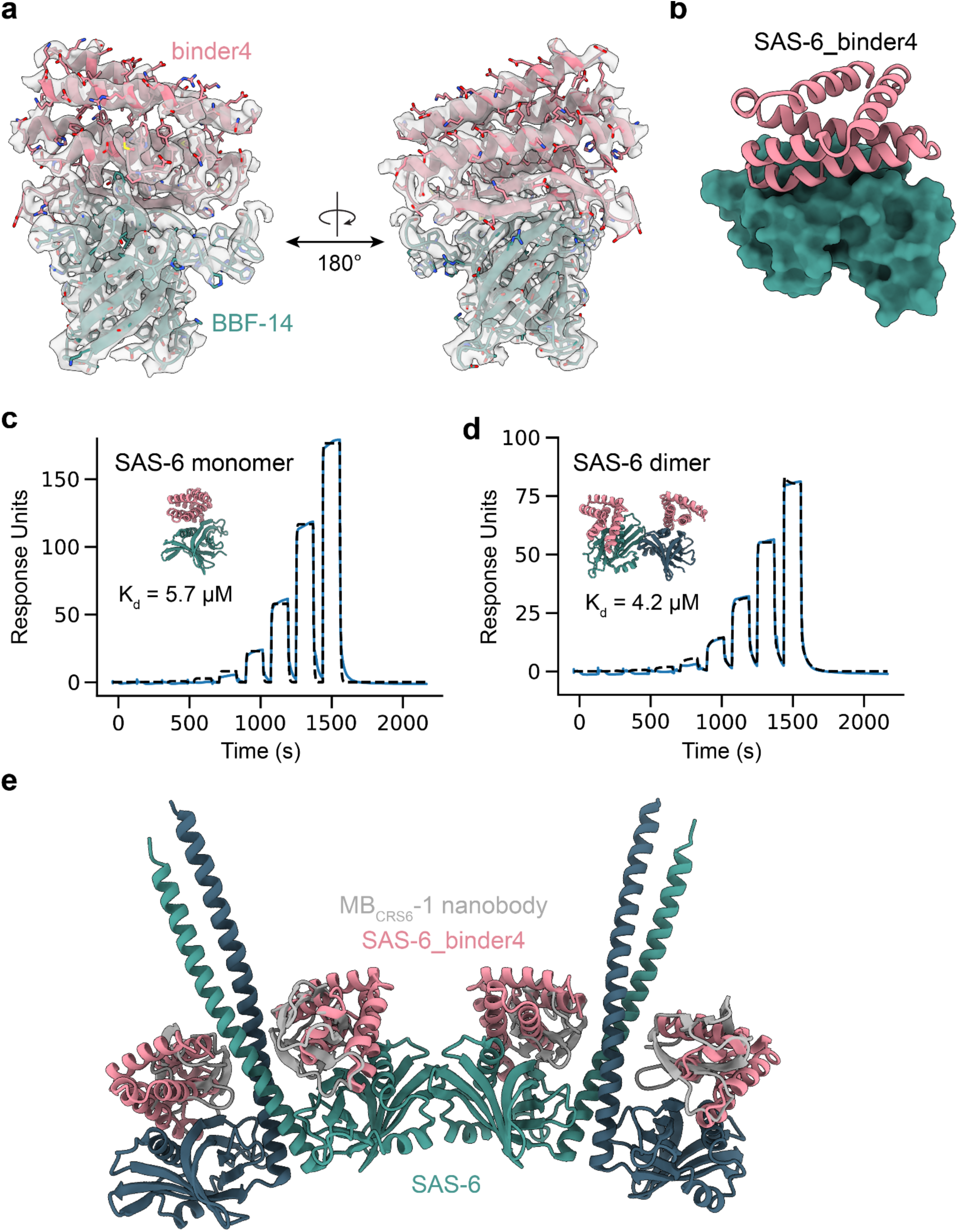
Design of SAS-6-targeting binders. **a,** Refined 2mFo − mFc electron density map of the BBF-14_binder4 complex rendered in gray and contoured at 1.0σ. The model complex refined against the map is shown as cartoon representation with BBF-14 colored green and binder4 in salmon. **b,** Design model of binder4 binding to the challenging structural protein target SAS-6. **c,** SPR binding traces of binder4 to CrSAS-6 monomeric form. **d,** SPR binding traces of binder4 to CrSAS-6 dimeric form. **e,** Structural model of the oligomeric form of SAS-6 with binder4 (salmon) overlaid with the previously characterized monobody MB_CRS6_-1^27^.

**Extended Data Fig. 6.**
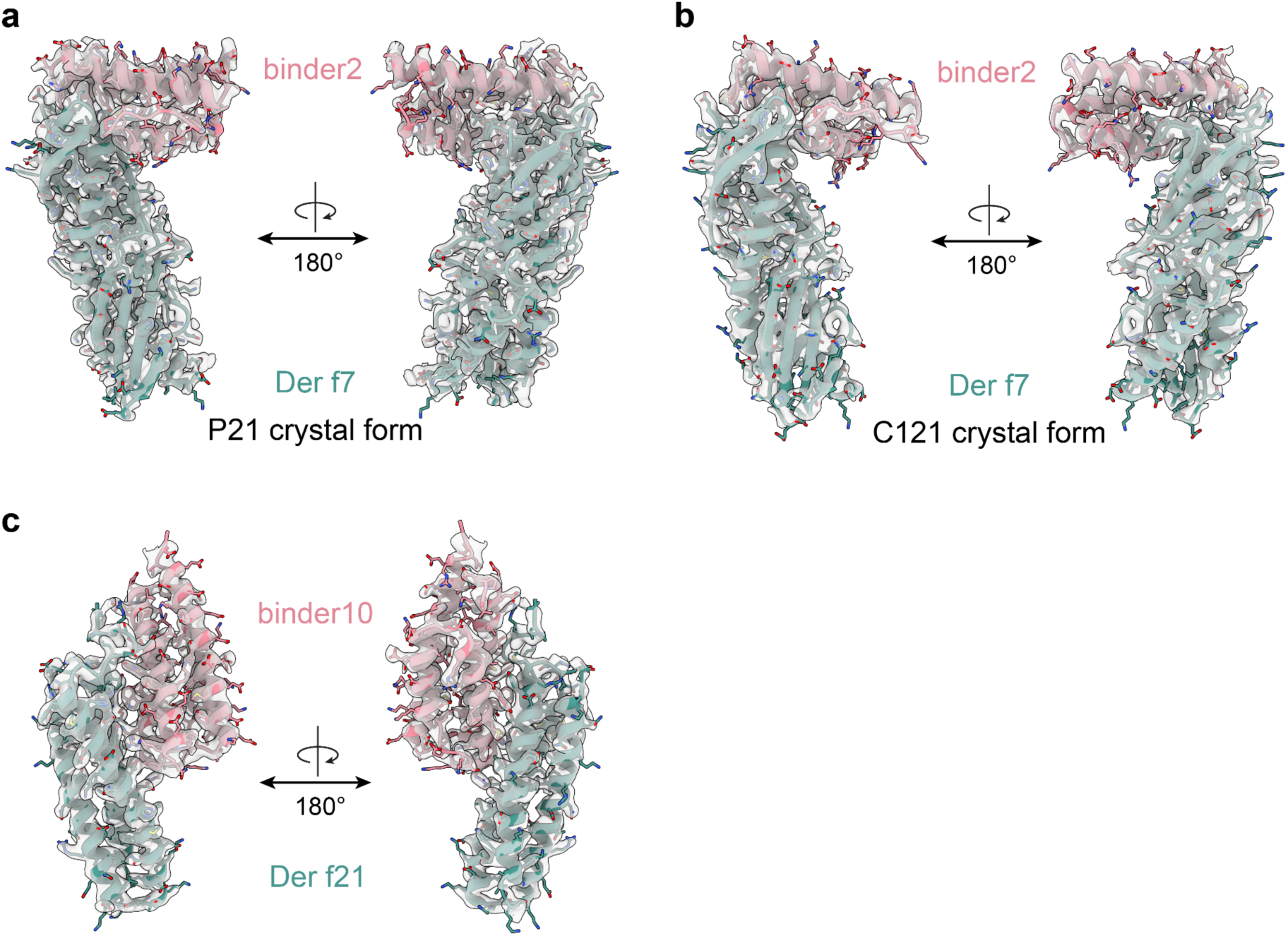
Crystallographic structures of dust mite allergens bound to designed binders. Refined 2mFo − mFc electron density map of the DerF7_b2 complex rendered in gray and contoured at 1.0σ for crystal form P21 in **a** and the crystal form C121 in **b**. The model complex refined against the map is shown as cartoon representation with Der f7 colored green and binder2 in salmon. **c,** Refined 2mFo − mFc electron density map of the DerF21_binder10 complex rendered in gray and contoured at 1.0σ. The model complex refined against the map is shown as cartoon representation with Der f21 colored green and binder10 in salmon.

**Extended Data Fig. 7.**
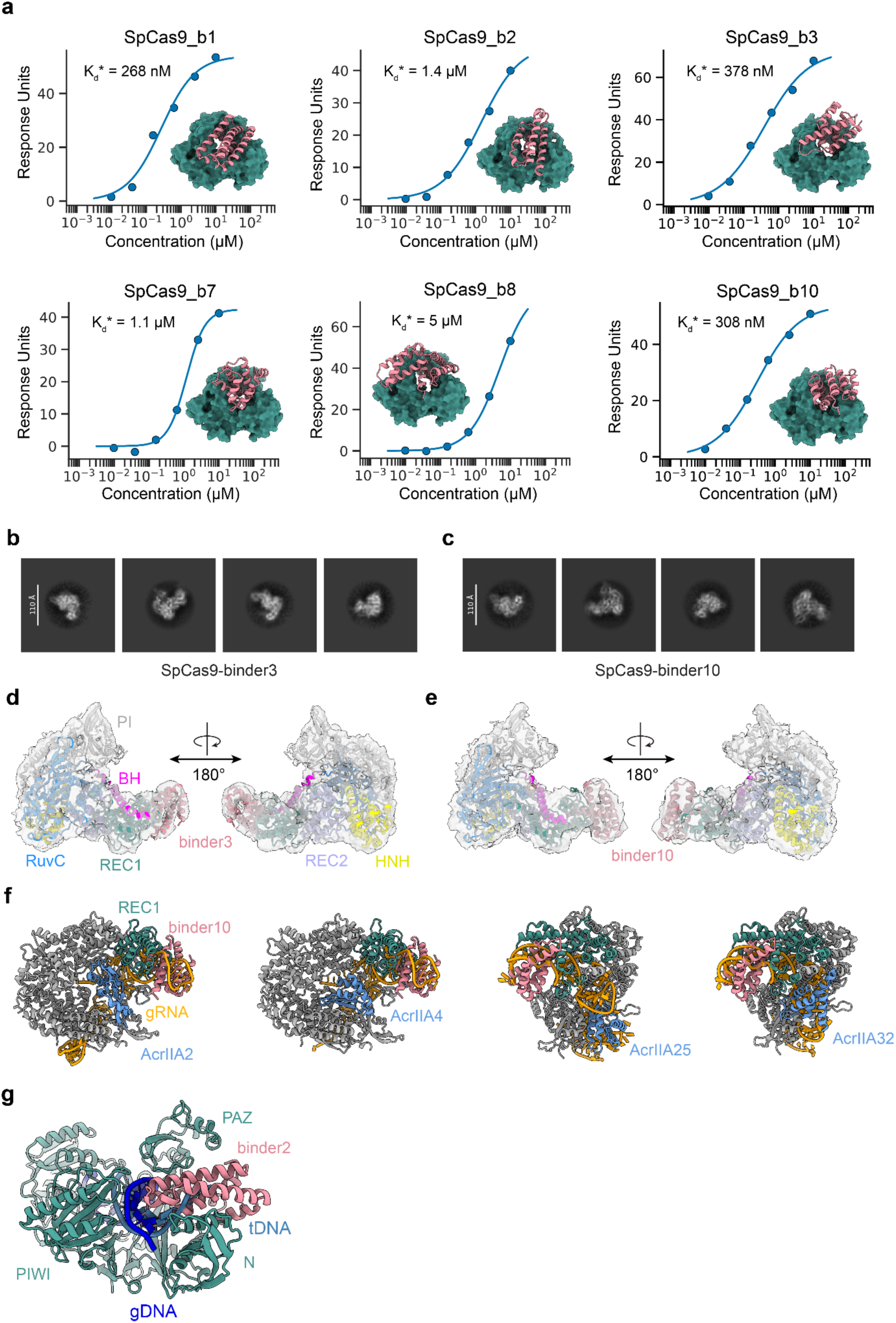
Biophysical and structural analysis of binders against nucleic acid-guided nucleases. **a,** SPR binding affinity fits for six tested SpCas9 binders. Representative 2D class averages of apo SpCas9 bound to **b,** binder3 and **c,** binder10. Predicted model of the apo conformation of SpCas9 with bound **d,** binder3 or **e,** binder10 docked into its respective cryoEM density. **f,** Experimental structures of known natural anti-CRISPRs (Acrs) bound to SpCas9. PDB codes 6IFO^43^, 5VW1^43,79^, 8YE6 and 8YE9^80^. **g,** Structural comparison of the binder2 overlaid with the target DNA-bound structure of CbAgo, indicating overlapping binding sites.

**Extended Data Fig. 8.**
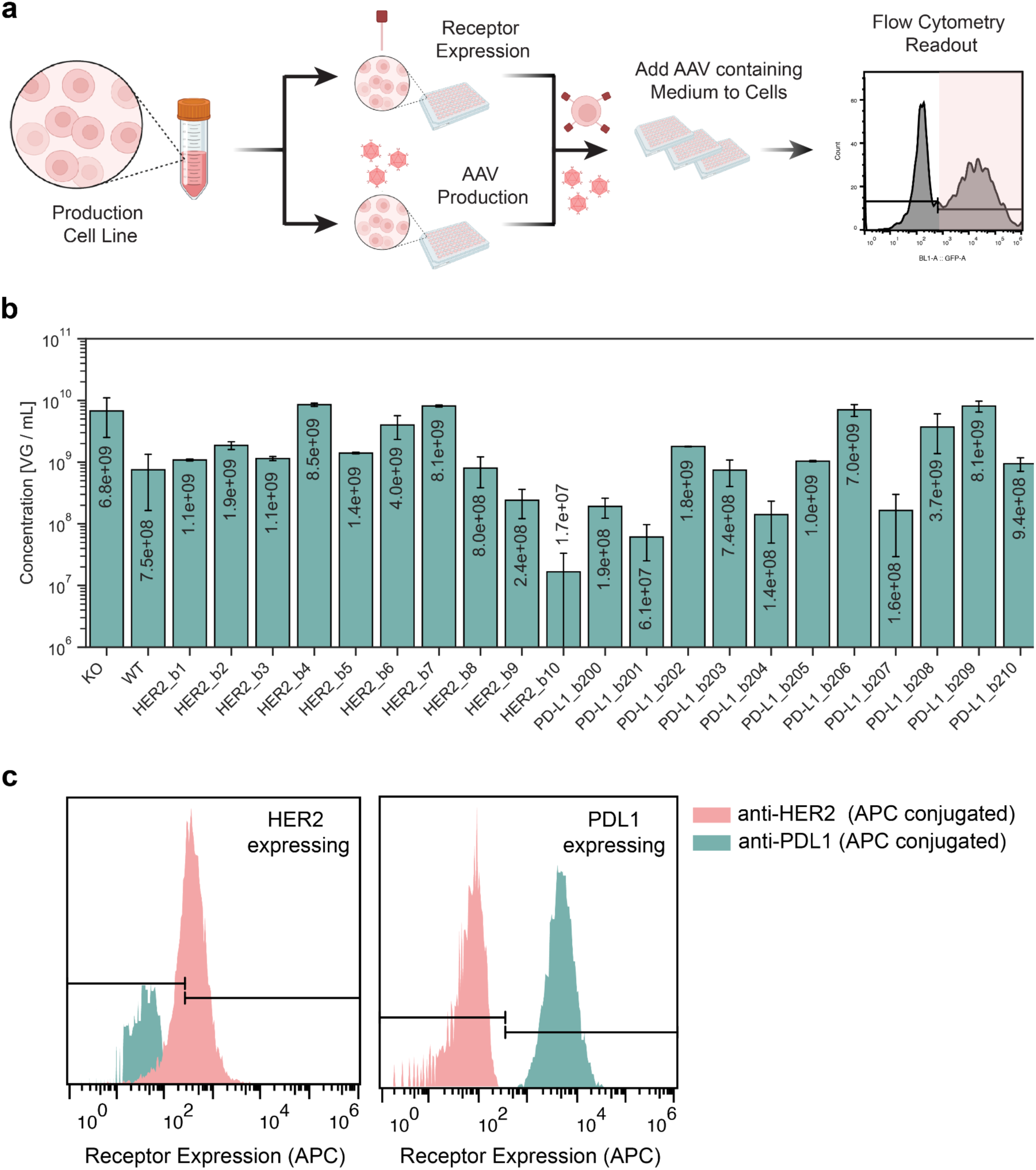
Screening of functional cell-type specific AAVs. **a,** Schematic illustrating the small-scale screening assay. Both the production cell line as well as the target cells overexpressing the target receptors are derived from the same parent cell line, allowing to directly transfer the supernatant of AAV-packaging cells onto the targeted cells for transduction. The right scatter plot illustrates the transduction signal measured by flow cytometry (GFP expression) **b,** Supernatant viral titers (log-scale) of the different AAV variants screened (Fig. 6c), as indicated. **c,** Receptor expression levels of the created stable cell lines for screening, stained by APC-conjugated antibodies.

**Extended Data Fig. 9.**
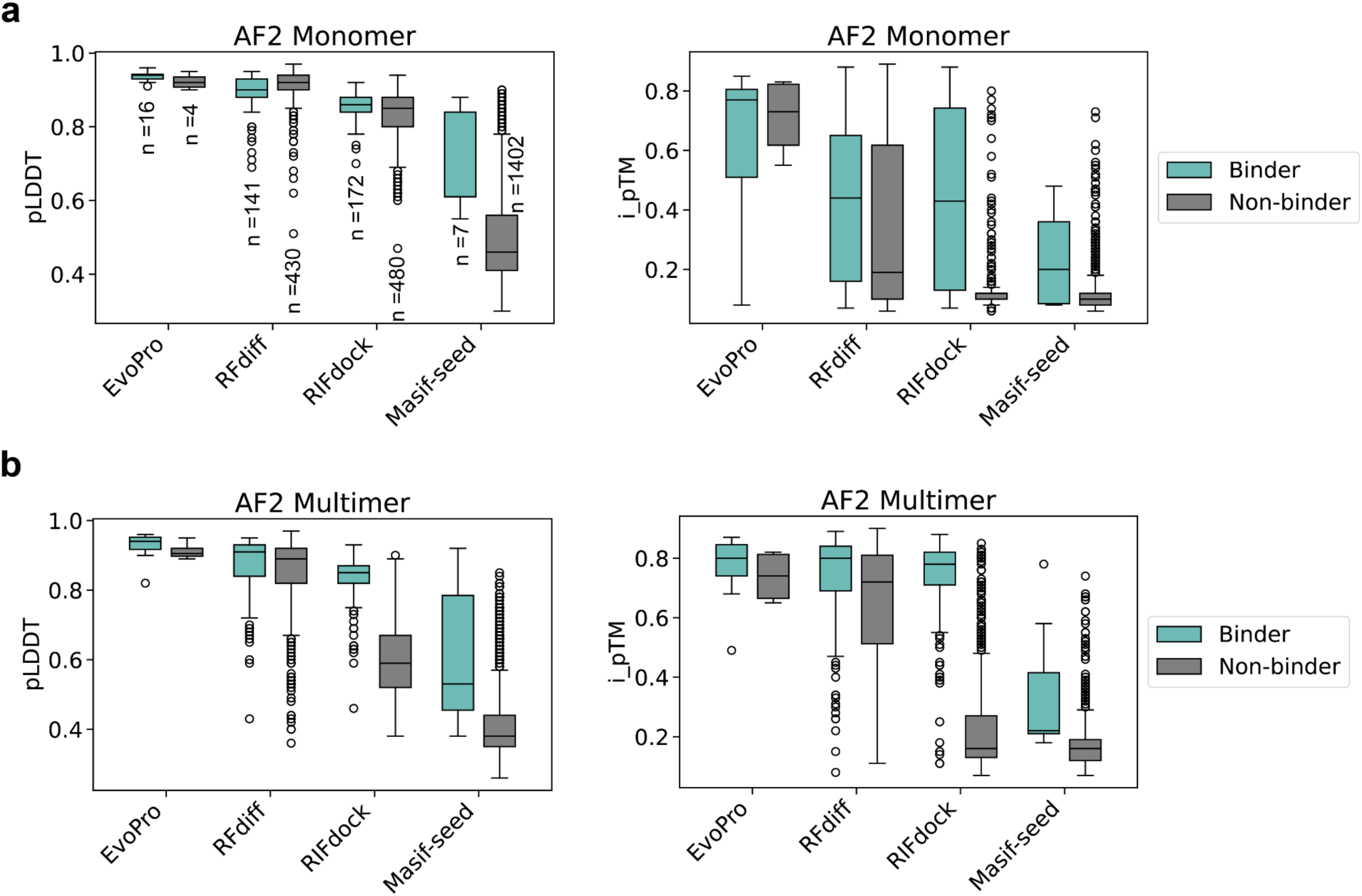
Comparing AlphaFold2 prediction accuracy across design pipelines. Experimentally validated binders and non-binders from previously published binder design pipelines have been repredicted using the BindCraft prediction pipeline with either AF2 **a,** monomer (default) or **b,** multimer models. Of note, EvoPro^56^ and RFdiff^10^ designs have been already prefiltered by AF2 monomer in their respective publications, and indicate the presence of false positives. RIFdock^11^ and Masif-seed^8,11^ designs were not prefiltered by AF2.

